# DiffBreed: Automatic differentiation enables efficient gradient-based optimization of breeding strategies

**DOI:** 10.1101/2024.11.29.625951

**Authors:** Kosuke Hamazaki, Hiroyoshi Iwata, Koji Tsuda

## Abstract

**Motivation:** Differentiable programming frameworks like PyTorch and JAX revolutionized biological modeling. A foremost merit is that multiple components programmed separately can be put together so that the parameters are jointly optimized. Despite its proven value in agricultural applications, existing breeding simulators are non-differentiable, hindering integration into general deep learning systems.

**Results:** In this paper, we present DiffBreed, a differentiable breeding simulator. Its performance was evaluated in gradient-based optimization of a progeny allocation strategy that maximizes the genetic gain. By utilizing gradient-based optimization, DiffBreed refined progeny allocation strategies, achieving superior genetic gains compared to a non-optimized equal allocation approach. These findings highlight DiffBreed’s capacity to properly calculate gradient information through automatic differentiation. With its innovative design, DiffBreed is expected to transform future breeding optimization by integrating into modern deep learning workflows.

**Availability:** The proposed utomatic differentiation framework was implemented as the Python module “DiffBreed”. This module and all the scripts used in this study, including the gradient-based optimization, are available from the “KosukeHamazaki/GORA-PT” repository on GitHub, https://github.com/KosukeHamazaki/GORA-PT. While the simulated datasets in the present study are available from the same repository, the optimized results by PyTorch were not shared in this repository due to the file sizes. Instead, all datasets, including the optimized ones, will be shared in the repository on Zenodo, https://doi.org/10.5281/zenodo.14046522.

## 1 Introduction

In recent years, machine learning and optimization techniques have transformed numerous fields by providing efficient solutions to complex problems. Similarly, in plant breeding, these techniques have become increasingly important for enhancing breeding strategies through more systematic approaches. Breeding simulators, such as AlphaSimR (Gaynor et al., 2021), PyBrOpS (Shrote and Thompson, 2024), and breedSimulatR (Diot, 2025), serve as valuable tools that approximate real breeding schemes and help optimize these strategies in virtual environments. These simulators implement selection and mating steps that generate new populations from previous ones, effectively mimicking the gradual improvement of genetic merit in breeding populations (Figure 1). Through iterative selection and mating processes, these simulators allow researchers to predict and evaluate the effectiveness of different breeding approaches without the time constraints of physical breeding cycles.

**Fig. 1:**
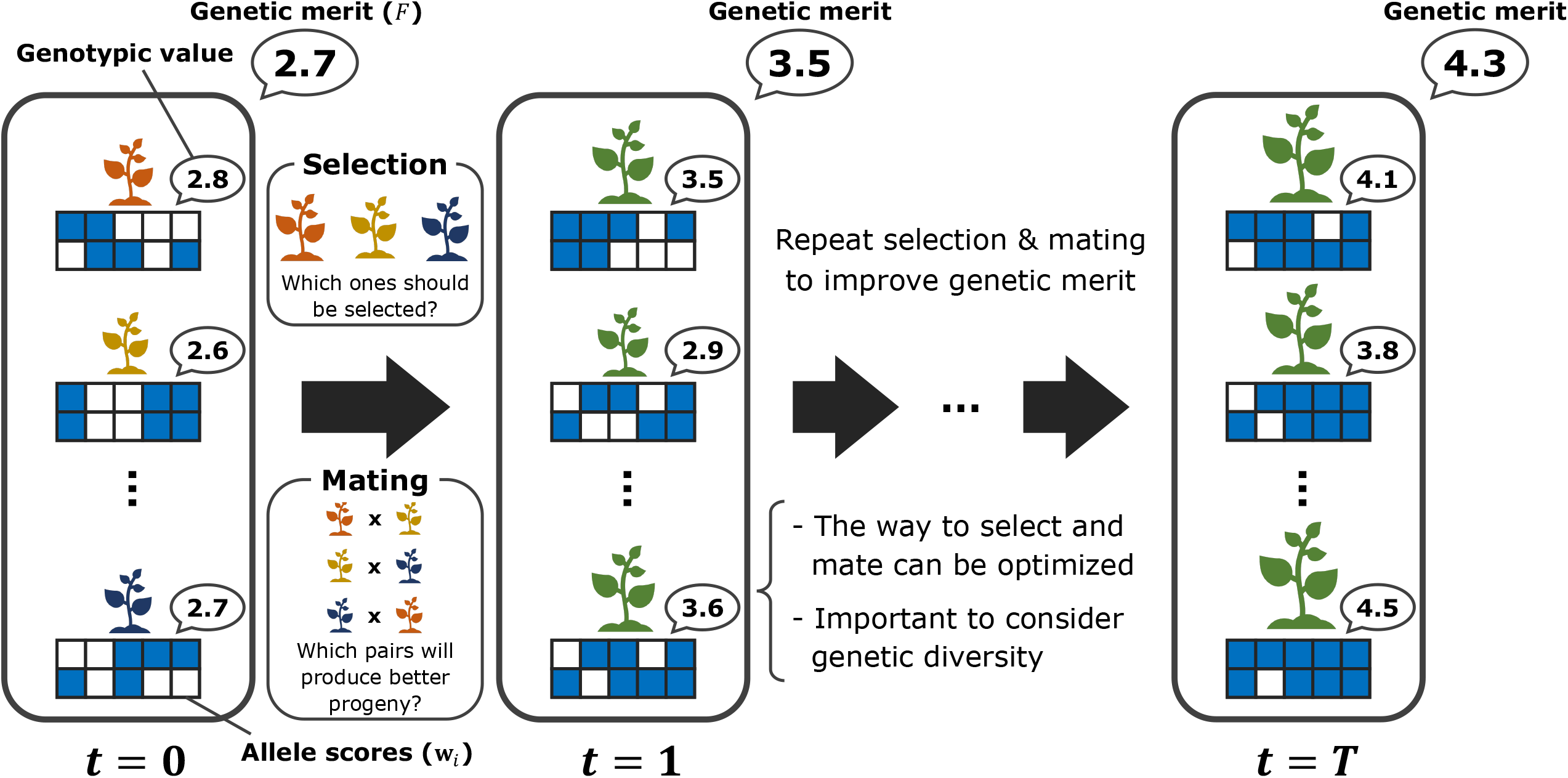
Breeding simulators in case of recurrent selection. Breeding simulators approximate breeding schemes that aim to improve the genetic merit of a population through iterative selection and mating processes.

Since genetic merit is directly influenced by selection and mating decisions, researchers have begun exploring various simulator-oriented optimization methods. However, this optimization faces a significant challenge: focusing solely on selecting superior individuals inevitably reduces genetic diversity over generations, causing long-term improvement to plateau. To overcome this problem, optimal contribution selection (Meuwissen, 1997) focuses on balancing genetic gain and diversity by optimizing individual contributions to the next generation. Another approach, look-ahead selection (Moeinizade et al., 2019) anticipates potential breeding outcomes to make better crossing decisions under simplified assumptions. Beyond these approaches, research has explored other optimization techniques, including branch-and-bound method (Hunter and McClosky, 2016), integer programming (Sakurai et al., 2024), Bayesian optimization (Diot and Iwata, 2022; Jannink et al., 2023), and reinforcement learning (Moeinizade et al., 2022), to determine appropriate breeding strategies and planning. These studies have been driven by various breeding simulators, such as AlphaSimR (Gaynor et al., 2021), PyBrOpS (Shrote and Thompson, 2024), and breedSimulatR (Diot, 2025), which have enabled researchers to evaluate specific breeding strategies. Hamazaki and Iwata (2024) advanced this area by optimizing progeny allocation strategies through a combination of future-oriented breeding simulations and black-box optimization. They conceptualized the breeding scheme as a black-box function, with parameters related to progeny allocation as inputs and the final genetic gains as outputs, and applied a black-box optimization technique called StoSOO to optimize it (Valko et al., 2013). Their method provided a flexible framework capable of handling complex breeding schemes without restrictive assumptions by allowing customizable breeding simulation settings.

However, traditional simulators used for optimizing breeding strategies are non-differentiable, which limits flexible optimization when managing large-scale breeding schemes or optimizing multiple breeding aspects simultaneously. Non-differentiable simulators cannot effectively leverage gradient-based optimization, resulting in slower convergence and reduced performance, while also limiting their versatility in handling complex computational models (Baydin et al., 2018). Automatic differentiation (AD) has recently emerged as a powerful computational method that can replace traditional non-differentiable simulators with differentiable alternatives (Rall, 1981; Griewank and Walther, 2008; Baydin et al., 2018). AD automatically computes derivatives of complex functions defined in computer programs, enabling rapid gradient calculations that significantly accelerate optimization processes. This technique has been successfully applied across diverse scientific domains, including machine learning (Baydin et al., 2018), physics (Callejo et al., 2014; Luchnikov et al., 2021), chemistry (Tamayo-Mendoza et al., 2018; Yoshikawa and Sumita, 2022), and biology (AlQuraishi and Sorger, 2021; Frank, 2022). Programming frameworks like TensorFlow (Martín Abadi et al., 2015), PyTorch (Paszke et al., 2017), and JAX (Bradbury et al., 2021) have facilitated this broad adoption by simplifying implementation, which has led to differentiable forms increasingly replacing traditional simulators.

Therefore, integrating AD into plant breeding optimization offers a promising solution to address the technological gap described above. While AD has shown promise in agricultural applications, such as crop growth modeling for biomass optimization (Lauvernet et al., 2012), it has not yet been applied specifically to breeding optimization. Thus, we focus on optimizing progeny allocation in breeding schemes based on simple recurrent selection, expanding on previous optimization frameworks by Hamazaki and Iwata (2024) to incorporate gradients derived from AD. Our approach treats the entire breeding scheme as a differentiable computational graph, enabling efficient optimization through gradient-based methods. This framework, DiffBreed, when successfully implemented, could significantly enhance breeding efficiency through the optimization of breeding strategies using AD. As discussed in Section 4, this approach ultimately aims to enable a more flexible and adaptive breeding framework that, in the future, has the potential to contribute to sustainable food production as we face global challenges like population growth and climate change (Capper, 2011).

## 2 Methods

### 2.1 Basic idea

#### 2.1.1 Differentiable genetic merit of current population

Let us consider a breeding population consisting of *N* individuals, each represented by a binary vector **w**_*i*_ ∈ {0, 1}^*M*^. Here, *M* = 2*m* where *m* is the number of markers, and **w**_*i*_ represents allele scores formed by vertically stacking two haplotypes. Now, each individual can be evaluated by a genotypic value defined as ***α***^⊤^**w**_*i*_, where ***α*** ∈ ℜ^*M*^ is a vector of corresponding effects created by duplicating marker effects. We consider the problem of sampling *n* individuals (indexed as *z*_1_, …, *z*_*n*_) from the population based on a breeding strategy represented by a parameter vector ***θ*** ∈ ℜ^*d*^, i.e., *z*_1_, …, *z*_*n*_ are sampled from a parametric discrete distribution *P* (*z* | ***θ***). Without loss of generality, *z*_1_, …, *z*_*n*_ are assumed to be sorted in the descending order of the corresponding genotypic value 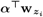. Then, when genetic merit of the population *F* is defined as the mean of genotypic values of top *K* individuals, it will be

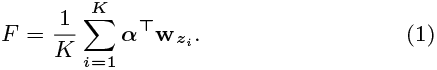

Genetic merit in this context refers to a case where selection exclusively targets superior individuals. Here, genetic gain can be defined as the difference in genetic merit between the initial population and the current population.

To evaluate how the parameter vector ***θ*** affects the genetic merit of the population *F*, we compute the derivative *∂F/∂****θ*** using the reparameterization trick (Jang et al., 2016; Maddison et al., 2017), which enables differentiable sampling from target distributions. Here, the selection of superior individuals is inherently a discreten and non-differentiable operation. Thus, by using the Gumbel-Softmax reparameterization trick, we replace this discrete selection with a continuous approximation. Specifically, when applying this trick, *y*_*ij*_ is sampled from the following Gumbel-Softmax function instead of directly sampling *z*_*i*_:

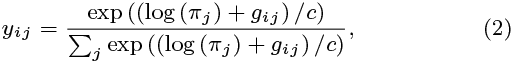

where *π*_*j*_ *∝ P* (*z* = *j* | ***θ***) represents the probability that an individual *j* will be selected when the parameter of the breeding strategy *θ* is determined, *g*_*i*1_, …, *g*_*iN*_ are i.i.d. samples drawn from Gumbel (0, 1), which enables stochastic sampling from categorical distributions, and *c* corresponds to the temperature parameter that balances discreteness and smoothness in the sampling process. Note that as *c* → 0, **y**_*i*_ = [*y*_*i*1_, …, *y*_*ij*_, …, *y*_*iN*_]^⊤^ becomes a one-hot encoded vector (a binary vector containing all zeroes except for a single element being one). Then, the sampled allele scores can be approximated as

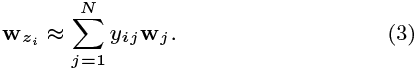

Since *F* is ultimately a linear function of *y*_*ij*_, we can apply the chain rule of calculus throughout the entire computation graph, which makes *F* differentiable with respect to ***θ***. Note that we introduced this subsubsection to better clarify the concept of differentiable merit. In this study, we specifically focused on optimizing the parameters related to the allocation strategy outlined in Section 2.1.2, rather than the selection strategy, as explained in Section 2.4.

#### 2.1.2 Differentiable genetic merit of next generation

Next, let us assume that progeny in the next generation is produced from the above *n* individuals in the current population. These *n* individuals undergo diallel crossing with selfing (all-to-all mating), creating *n*(*n* + 1)*/*2 mating pairs. Here, following Hamazaki and Iwata (2024), this subsubsection aims to determine how many progeny to allocate to each mating pair based on some “goodness” of each pair. However, this allocation process is a discrete and non-differentiable operation, which requires a continuous approximation, similar to the case in Section 2.1.1. We reframed this challenge by focusing on which mating pair to select as parents for each progeny in the next generation. We then approximated this pair selection process using differentiable sampling with the Gumbel-Softmax reparameterization trick, similar to the approach in Section 2.1.1.

To implement this approximation and obtain the allele score vector for a progeny 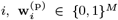, where (p) represents the genomic information of the “progeny”, we need to generate one offspring from each possible mating pair, then weight these allele score vectors using the *y*_*ij*_ in equation (2). An allele score vector of an offspring *i* from mating pair *k* ∈ {1, …, *n*(*n* + 1)*/*2}, 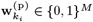 is determined by a function that takes allele score vectors of parents and a vector of random variables 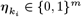. Here, 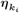 acts as the source of genetic variation between siblings from the same mating pair. Therefore, for the selection of mating pairs, equation (3) can be rewritten as follows:

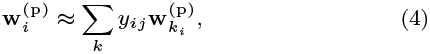

where the summation is taken over each mating pair rather than over each individual. To define the categorical distribution from which we aim to sample *y*_*ij*_, we also introduce a feature vector ***ω***_*k*_ ∈ ℜ^*d*^ representing “goodness” metrics for the mating pair *k*, which is calculated from the allele score vectors of the parents, with details described in Section 2.3. By utilizing the feature vector ***ω***_*k*_ and the parameter vector ***θ***, we define the parametric discrete distribution as follows:

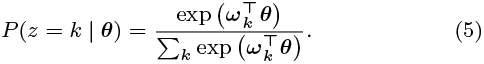

Note that in this subsubsection, *z* refers to the selected mating pairs rather than the selected individuals. Here, ***θ*** can be interpreted as weights for goodness metrics that determine the progeny allocation to each mating pair, making it the key parameter to be optimized in this study. Then, a sampled allele score vector of progeny 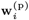 can be approximated by applying equations (2), (4), and (5). Consider that *N* individuals are produced in this manner for the next generation, and let us denote their differentiable genetic merit as *F* (***θ*, g, *η***), which can be computed by replacing **w**_*i*_ with 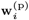 in equation (1). Unlike standard gradient-based optimization that targets deterministic functions, our genetic merit function *F* is stochastic because it incorporates two types of random factors: **g** and ***η***. First, **g** = {*g*_*ik*_} contains samples from the Gumbel (0, 1) distribution, formed by row-wise flattening of a *N × n* (*n* + 1) */*2 matrix. This introduces stochasticity when we approximate the breeding process in a continuous, differentiable form using Gumbel-Softmax reparameterization. Second, 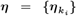 is a vector formed by flattening a *N × n* (*n* + 1) */*2 *× m* array, which causes the randomness in Mendelian segregation during gamete generation. Thus, our objective is to determine ***θ*** that optimizes

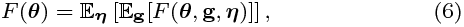

where 𝔼_**g**_ and 𝔼_***η***_ represent expectations with **g** and ***η***, respectively.

Using AD techniques provided in PyTorch (Paszke et al., 2017, 2019), we can easily extend this differentiable merit across multiple generations by repeating the above selection and mating process multiple times, allowing us to evaluate the derivatives for genetic merit at any generation *t* ∈ {0, …, *T* }, where *T* is the final generation (Supplementary Figure 1). When optimizing breeding schemes with multiple generations, each generation *τ* ∈ {0, …, *T* − 1} has its own distinct parameter vector ***θ***^(*τ*)^. This means that the genetic merit in equation (6) becomes a function of parameter vectors over multiple generations: *F* ({***θ***^(*τ*)^ }). As described above, by leveraging PyTorch’s computational graph architecture, we make the entire breeding process differentiable across all generations. Then, we can compute partial derivatives *∂F/∂****θ***^(*τ*)^ via AD. Finally, we used stochastic gradient descent (SGD) (Robbins and Monro, 1951) with these derivatives to optimize the allocation parameter ***θ***^(*τ*)^ (Supplementary Figure 2). The details of the breeding scheme with multiple generations and the evaluation of the optimized strategy are further elaborated in Section 2.4.

### 2.2 Formulation of progeny generation

In this subsection, we describe the details of how an allele score vector of the progeny 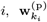, is generated from its parents named *k*_M_ and *k*_P_. When progeny are generated, gametes are independently produced from the parents *k*_M_ and *k*_P_ before being combined to form progeny. In the following, since the first *m* elements in 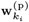 correspond to a gamete vector obtained from *k*_M_ and the latter *m* elements correspond to that from *k*_P_, we focus on the gamete generation process from *k*_M_, as the procedure is identical for both.

In this process, a recombination event occurs in which the maternal and paternal haplotypes of a parent are mixed together, and subsequently, one of these mixed haplotypes is generated as a gamete (Supplementary Figure 3). Here, whether information exchanges occur between markers *j* and *j*^*′*^ (1 ≤ *j, j*^*′*^ ≤ *m*) is determined by sampling based on a recombination rate 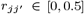. We note that 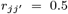 if two markers *j* and *j*^*′*^ are located on different chromosomes. Let us denote 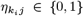 as a sample from a Bernoulli distribution with probability *r*_*j*−1*j*_ where we virtually set *r*_01_ = 0.5 to randomly select either maternal or paternal haplotype with equal probability. Here, the cumulative summation of 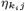 represents the total number of crossover events. Since an even number of crossover events restores the original haplotype, we can express an allele score at marker *j* for gamete *i* from mating pair *k* as follows.

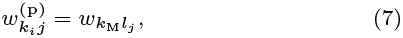

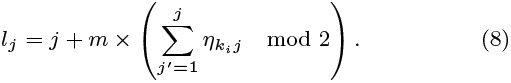

These equations demonstrate that the allele score of a gamete is directly inherited from one of the parent’s haplotypes at each marker, and which specific haplotype is inherited depends on the number of crossover events that have occurred up to that point in the chromosome. We implemented this progeny generation method using PyTorch.

### 2.3 Goodness metrics employed for progeny allocation

As the candidates for the feature vector of “goodness” metrics ***ω***_*k*_ ∈ ℜ^*d*^ in the allocation step, we used a selection criterion called weighted breeding value (WBV) (Goddard, 2009; Jannink, 2010) and the expected genetic variance of progeny (GVP) for each mating pair, i.e., *d* = 2. WBV is a criterion that enables genetic improvement while maintaining genetic diversity in a breeding population by placing greater emphasis on rare alleles, and is considered advantageous for long-term breeding schemes. In this study, WBVs are calculated for both parents *k*_M_ and *k*_P_, and their mean serves as the first element of ***ω***_*k*_ as follows (Jannink, 2010).

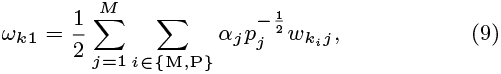

where *p*_*j*_ is a minor allele frequency at marker *j*.

We also examine GVP as the genetic diversity of gametes produced after the self-fertilization of progeny from each mating pair to assess the genetic distance between parents. Here, this GVP can be theoretically derived as in the following equation (10), based on the idea proposed by previous studies (Lehermeier et al., 2017; Allier et al., 2019a,b), serving as the second element of ***ω***_*k*_.

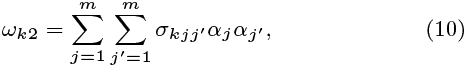

where 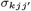 is a covariance between markers *j* and *j*^*′*^ caused by the segregation of progeny derived from mating pair *k*. The detailed deviation of 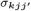 is provided in Allier et al. (2019b), but can be computed based on the recombination rate 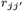 and allele score vectors of both parents (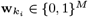 for *i* ∈ {M, P}) as follows:

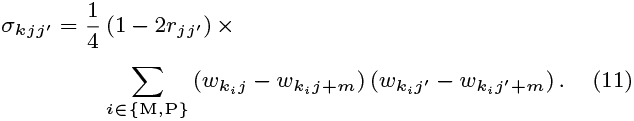

### 2.4 Evaluation of the proposed framework

As outlined by Hamazaki and Iwata (2024), we simulated a breeding scheme over *T* = 4 generations using a progeny allocation strategy proposed by an “AI breeder” (Supplementary Figure 2). In this study, we did not implement the method described in Section 2.1.1 at the selection stage. Instead, we used predetermined selection strategies described below and focused primarily on optimizing the parameters related to the progeny allocation strategy ***θ***^(*τ*)^ explained in Section 2.1.2 for simplicity. Here, breeders were assumed to obtain the optimal allocation parameters 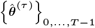 by providing the AI breeder with initial population information, including allele scores, marker effects, and recombination rates.

To optimize ***θ***^(*τ*)^, the AI breeder assessed allocation strategies by measuring the final genetic gain—defined as the relative genetic merit of the final population in equation (1) of the top *K* = 5 individuals compared to the initial population. As discussed in Section 2.2, since the genetic gain includes two random factors, the AI breeder needs to maximize the expected final genetic gain given in equation (6). Here, the expectations with respect to the random variables **g** and ***η*** in equation (6) were handled by taking the empirical mean of the final genetic gains for 100 breeding simulations based on each ***θ***^(*τ*)^. Although computationally intensive, running 100 simulations was essential for accurately approximating the expected value of this stochastic breeding process, as gradient calculations based on a single simulation would produce prohibitively noisy and unreliable estimates. Then, after repeating the function evaluations of *F* 200 times in SGD (Robbins and Monro, 1951) while computing gradient information through AD approaches (Paszke et al., 2017) as described in Section 2.1.2, 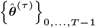 was passed from the AI breeder to the actual breeders.

In this study, to validate whether AD could contribute adequately to the gradient-based optimization of breeding schemes, we performed simulation studies as a proof of concept. We first simulated the initial breeding populations with a population size of *N* = 250, assuming a virtual diploid crop with ten chromosomes. Here, we maintained a constant population size of *N* = 250 throughout the entire breeding scheme in this study. Then, we prepared two different scenarios concerning the number of quantitative trait loci (QTLs): *m* = 30 (Scenario 1) or *m* = 500 (Scenario 2) in total. For simplicity and due to computational limitations, we also assumed true QTL positions and effects were known when determining breeding strategies. The details of simulating the initial population and QTLs are described in Supplementary Section S1.

In each scheme, we prepared the following three methods to select *n* individuals as parent candidates for mating based on the order of increasing WBV (Jannink, 2010), with details given in Supplementary Section S2 (Supplementary Figure 4).

1. SI1: Selected *n* = 15 genotypes from the current population.
2. SI2: First performed hierarchical clustering based on marker genotypes, creating 25 clusters. Then, it selected the only top genotype from each cluster, resulting in *n* = 25 genotypes.
3. SI3: Selected *n* = 50 genotypes from the current population.

Then, across these two scenarios featuring different QTL numbers and three selection strategies with different intensities, we compared the following four allocation strategies through 10,000 breeding simulations: two versions of the gradient-based optimized resource allocation (GORA1 and GORA2) based on DiffBreed, the black-box-based optimized resource allocation (ORA) by Hamazaki and Iwata (2024), and the non-optimized equal allocation strategies. The non-optimized equal allocation strategy refers to a simple baseline approach where we distribute the number of progenies equally across all mating pairs, which corresponds to ***θ***^(*τ*)^ = **0** in the proposed strategy. Here, the two versions of GORA differ in the initial weights corresponding to GVP (*ω*_*k*2_), i.e., the initial weight of GORA1 was larger than that of GORA2, with details given in Supplementary Section S3. We note that the domain of definition of ***θ***^(*τ*)^ was defined as ***θ***^(*τ*)^ ∈ [0.5, 3.5]^*d*^ using the inverse logit transformation of ***ν***^(*τ*)^ ∈ [−∞, ∞]^*d*^ for both GORAs and ORA. We implemented this inverse logit transformation to prevent parameters from converging to unreasonable values, while enabling SGD to solve the problem as an unconstrained optimization task. Also, the optimized parameters in ORA were obtained through 20,000 function evaluations, using 50 breeding schemes to calculate the expectation with respect to ***η***.

Throughout the study, R version 4.4.1 (R Core Team, 2024) was used to simulate the marker genotype of the initial breeding population, while Python version 3.12.2 (Van Rossum and Drake, 2009) with PyTorch version 2.3.1 (Paszke et al., 2017, 2019) was used to implement DiffBreed and conduct breeding optimization.

## 3 Results

### 3.1 Evaluation of convergence conditions

We first verified whether gradient-based optimization with AD could successfully optimize the allocation strategy by monitoring changes in function values across two scenarios under three selection strategies (Supplementary Figure 5).

The results showed that the function values generally improved with increasing SGD epochs, with the exception of Scenario 1 under SI1 (Supplementary Figure 5). Notably, allocation strategies under SI2 and SI3 were effectively optimized, with the algorithm converging to certain levels within 100 epochs (Supplementary Figure 5 C-F). These findings indicate that gradients derived from AD facilitated an efficient search for optimal parameters.

### 3.2 Genetic gains over four generations

We compared the four allocation strategies (GORA1, GORA2, ORA, and EQ) by analyzing the genetic gains over four generations across two scenarios under three selection strategies (Figure 2).

**Fig. 2:**
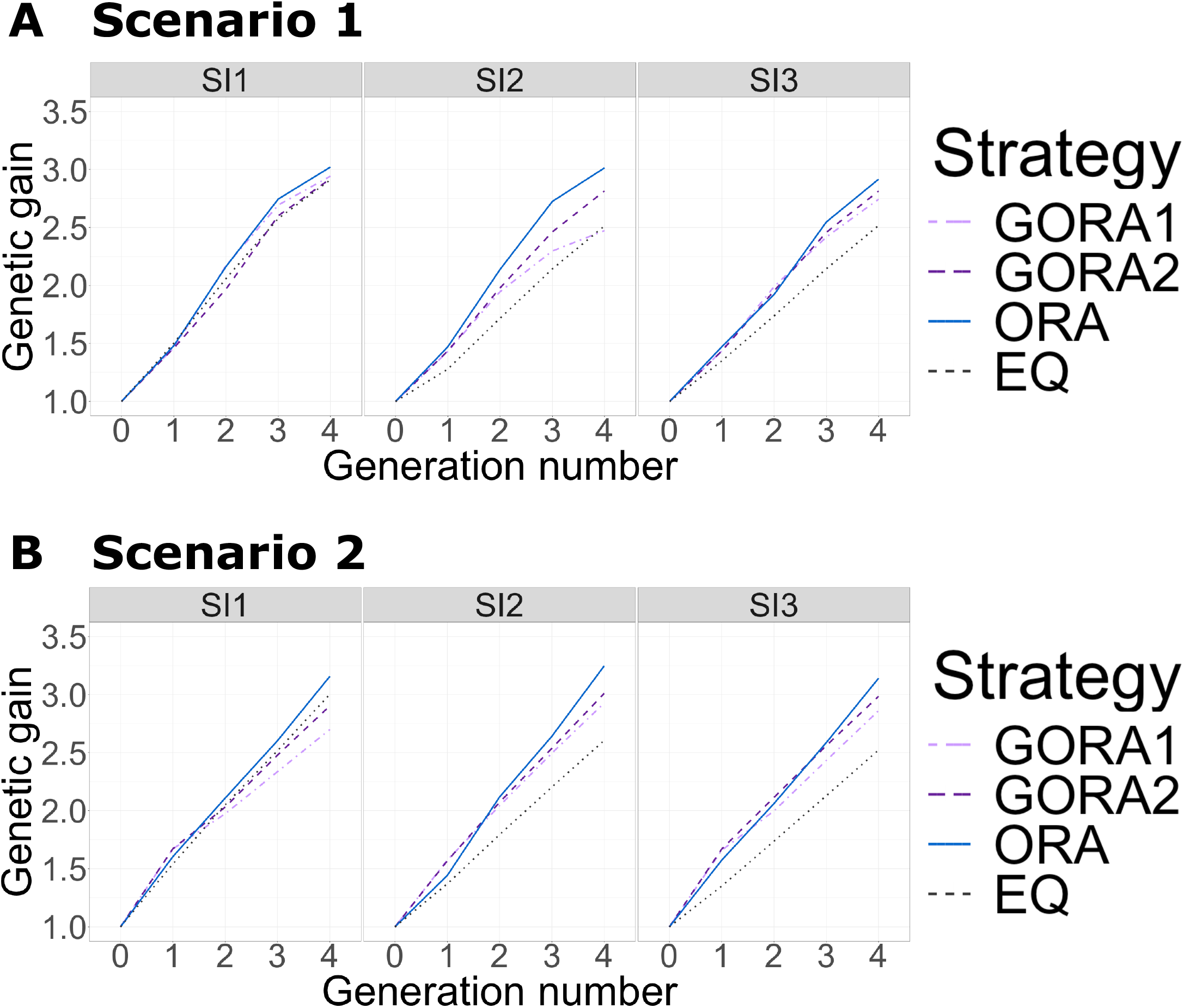
Change in the genetic gains over four generations under different selection intensities. The horizontal and vertical axes represent the number of generations and the genetic gains, respectively. We compared the four allocation strategies under different selection intensities (SI1 – SI3): GORA1: gradient-based optimized allocation with larger initial weights on GVP (light purple dash-dotted), GORA2: gradient-based optimized allocation with smaller initial weights on GVP (purple dashed), ORA: black-box-based optimized allocation (blue solid), EQ: equal allocation (black dotted). (A) Scenario 1. (B) Scenario 2.

In the final generation, *t* = 4, ORA demonstrated the highest genetic gain for both scenarios (Figure 2). Under strong selection intensity (SI1), there was minimal difference between the proposed and equal allocation strategies. However, under weaker intensities (SI2 and SI3), the proposed GORA strategies outperformed the non-optimized EQ strategy in both scenarios. The effectiveness of optimized allocation (ORA and GORA) was particularly pronounced when the trait was controlled by many QTLs, as in Scenario 2. Comparing the two proposed strategies, GORA2 generally outperformed GORA1. This suggests that the initial values significantly influenced the gradient-based optimization.

For Scenario 2, although ORA initially showed lower genetic gains than GORAs in *t* = 1, from *t* = 2, ORA overtook GORAs. This indicates that the black-box-based strategy successfully fine-tuned the balance between the weights on WBV and GVP.

### 3.3 Genetic gains across different simulation repetitions

To better understand the nuance of each strategy, we analyzed the cumulative distribution functions (CDFs) of genetic gain in the final generation (*t* = 4) for each allocation strategy, using 10,000 simulation repetitions. Supplementary Figure 6 displays these results, with the horizontal axis showing the final genetic gain of each breeding simulation and the vertical axis indicating the percentile of simulation repetitions. The 1st and 99th percentiles represent the worst and best performances among the 10,000 repetitions, respectively. The effectiveness of each strategy is demonstrated by its CDF curve shifting rightward and downward on the graph—the further this shift, the better the strategy’s overall performance.

Once again, ORA consistently showed the best performance (Supplementary Figure 6). Among the proposed strategies, GORA2 outperformed GORA1 in all cases and EQ under low selection intensities (SI2 and SI3). In Scenario 2, the CDF curves of the optimized strategies (GORA and ORA) exhibited shallower slopes compared to Scenario 1, suggesting that a larger number of QTLs led to greater variations among simulation repetitions. Particularly in Scenario 1, GORA2 displayed steeper slopes than ORA, indicating variable performance: at its best, it matched the ORA’s performance, but at its worst, it produced inferior results.

### 3.4 Genetic gains across different phenotype simulations

We further evaluated the final genetic gains using ten replications of the phenotype simulation to verify whether the gradient-based optimized strategies outperformed the equal allocation strategy for different target traits with varying QTL positions and effects (Figure 3). When simulating phenotypes for different traits, we altered QTL positions and their effects while keeping the number of QTLs and the distribution of QTL effects constant. Here, we chose GORA2 to represent our proposed approach, as it consistently outperformed GORA1 in all cases. To reduce computational time while ensuring that the function value reached a satisfactory level, we determined the optimized parameters of GORA after 100 function evaluations and ORA after 5,000, based on the convergence assessment in Section 3.1. Full convergence was not necessary, as our goal was simply to demonstrate that our algorithms could improve genetic gain compared to the non-optimized one.

**Fig. 3:**
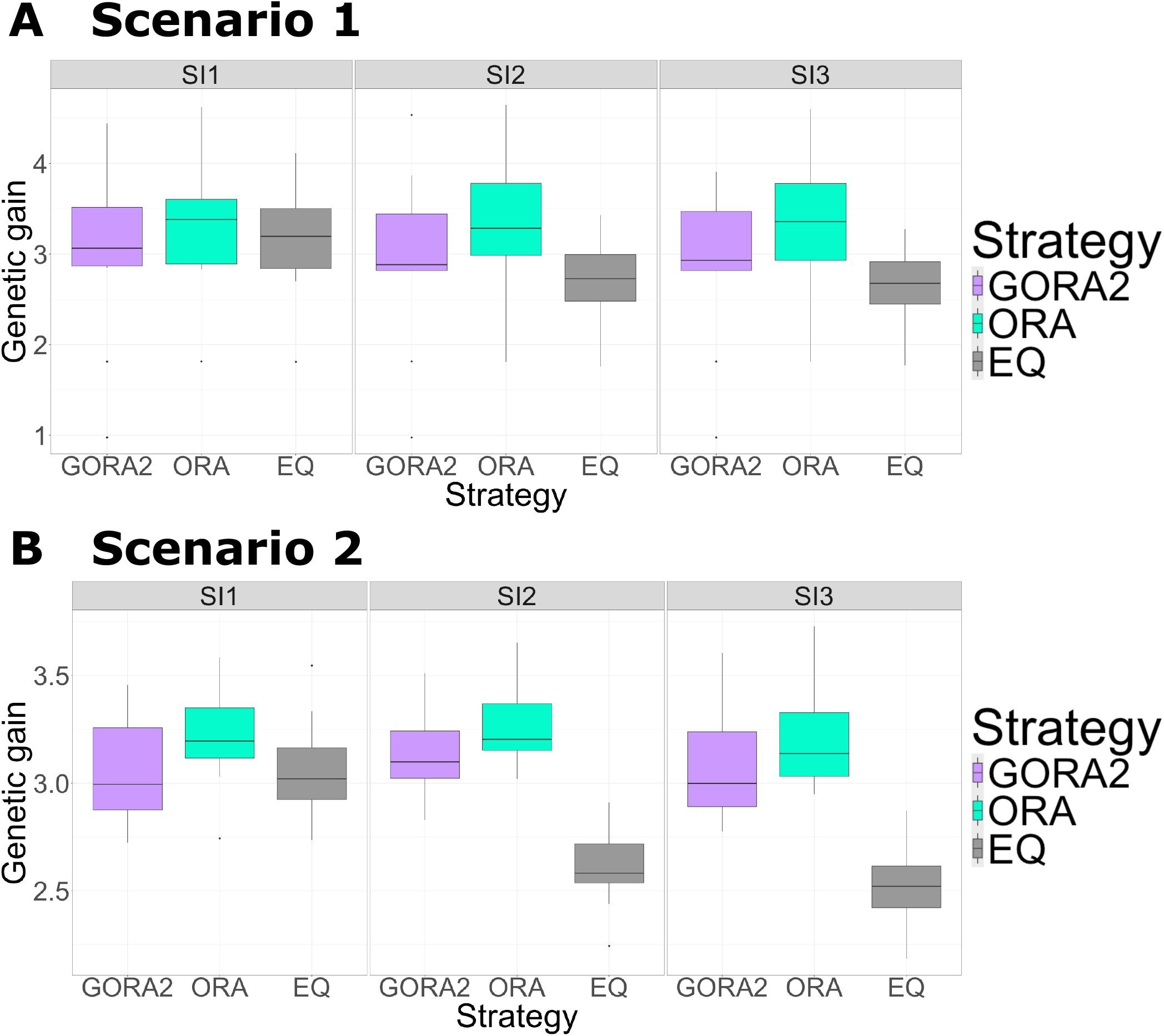
Genetic gains in the final generation using ten replications for the phenotype simulation under different selection intensities. Boxplots of the genetic gains in the final generation using ten replications for the phenotype simulation. The horizontal and vertical axes represent the different allocation strategies and the final genetic gains, respectively. We compared the three allocation strategies under different selection intensities (SI1 – SI3): GORA2 (light purple), ORA (light blue), and EQ (grey), with details given in Figure 2. The weighting parameters were optimized based on 100 and 5,000 function evaluations for GORA2 and ORA, respectively. (A) Scenario 1. (B) Scenario 2.

ORA consistently showed the highest genetic gains, even when QTL positions and effects were altered (Figure 3). In addition, GORA outperformed EQ under SI2 and SI3 while showing little difference under SI1. Once again, the performance gap between optimized (ORA and GORA) and non-optimized (EQ) strategies was more pronounced under SI2 and SI3 in Scenario 2.

We evaluated the improvement rate in final gain for GORA2 and ORA compared to EQ using ten phenotype replications, which showed a similar trend between GORA and ORA (Supplementary Figure 7). Notably, GORA failed to optimize and underperformed EQ in only one replication for Scenario 1. In all other replications, the genetic gains of GORA were comparable to or higher than EQ, with remarkable improvements of 15.5 − 29.8% over EQ under SI2 and SI3 within just four generations.

### 3.5 Genetic diversity over four generations

Next, we compared the GORA, ORA, and EQ strategies in genetic diversity across generations for two scenarios under three selection strategies (Figure 4). The genetic diversity of a breeding population in each generation was evaluated based on the genetic variance of the true genotypic values in the population.

**Fig. 4:**
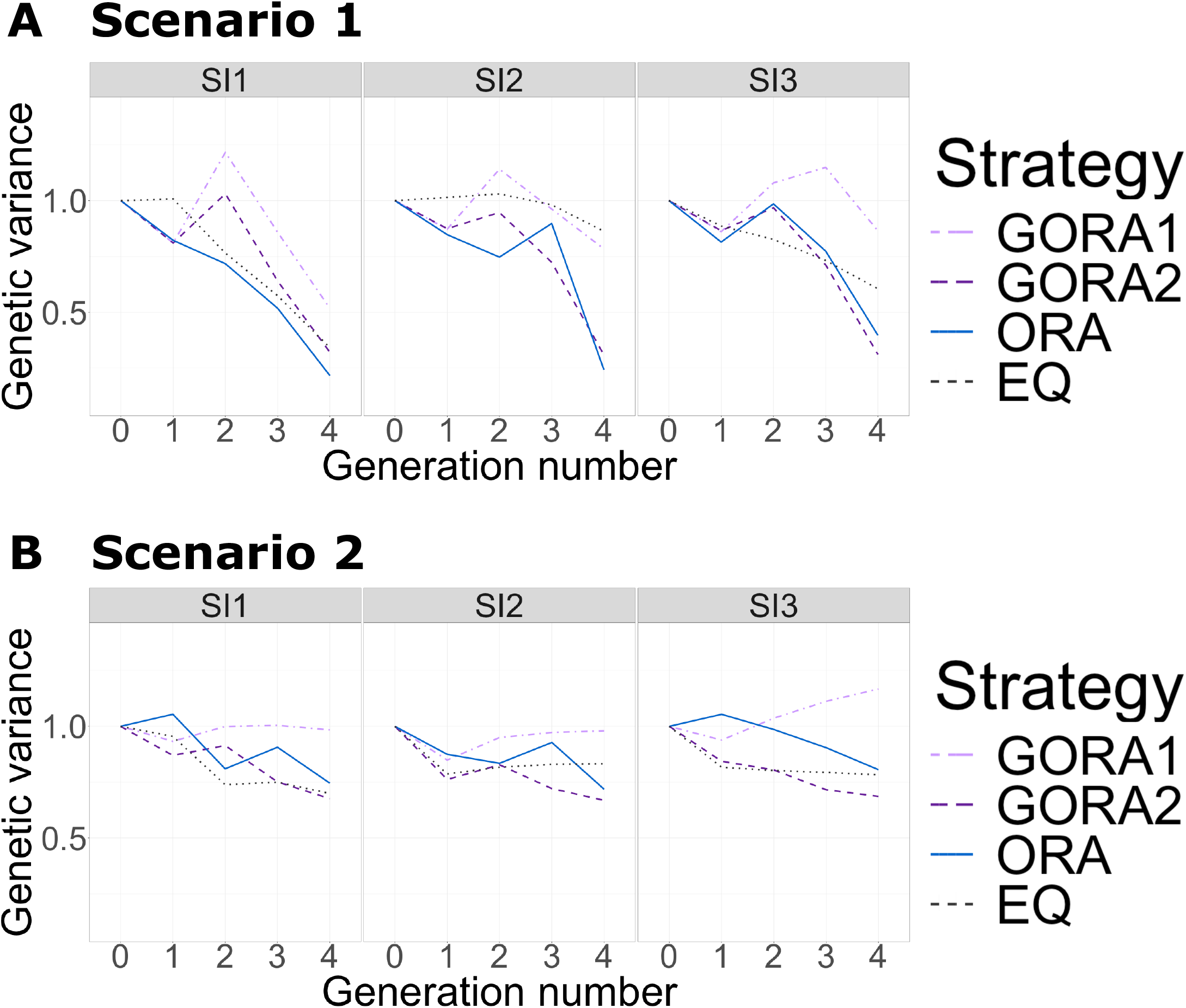
Change in the genetic variances over four generations under different selection intensities. The horizontal and vertical axes represent the number of generations and the genetic variances in the breeding population, respectively. We compared the four allocation strategies under different selection intensities (SI1 – SI3), with details of the abbreviations given in Figure 2. (A) Scenario 1. (B) Scenario 2.

First, in Scenario 1, GORA2 and ORA showed a rapid reduction in genetic diversity, while GORA1 relatively maintained the genetic variance throughout the scheme under all selection intensities (Figure 4A). In Scenario 2, the loss of diversity was suppressed compared to that in Scenario 1, and GORA2 showed the largest decrease in genetic diversity (Figure 4B). The genetic diversity of GORA1 in *t* = 4 was equal to or larger than that in *t* = 0. These results suggest that initial GVP weights in the allocation step largely impacted the genetic diversity of GORAs.

### 3.6 Optimized weighting parameters for each strategy

After 200 function evaluations for the GORAs and 20,000 for the ORA, we obtained optimized weighting parameters ***θ***^(*τ*)^ for two scenarios under three selection strategies (Table 1 and Supplementary Tables 1 - 5). Here, a higher value of ***θ***^(*τ*)^ indicates a greater emphasis on the corresponding goodness metric.

**Table 1.** Optimized weighting parameters *θ*^(*τ*)^ across generations *τ* = 0, …, *T* − 1 for Scenario 1 under SI1.

| $\tau$ | GORA1 | | GORA2 | | ORA | |
| --- | --- | --- | --- | --- | --- | --- |
|  | WBV | GVP | WBV | GVP | WBV | GVP |
| **0** | 2.63 | 0.38 | 2.93 | 0.20 | 1.50 | 0.17 |
| **1** | 2.09 | 1.92 | 2.13 | 0.29 | 1.50 | 0.17 |
| **2** | 2.26 | 1.93 | 2.33 | 0.04 | 2.83 | 0.17 |
| **3** | 2.22 | 1.99 | 2.39 | -0.09 | 2.83 | 0.17 |

Examining the weighting parameters for Scenario 1 under SI1 (Table 1), we find that GORA1 and GORA2 assigned higher weights to WBV than ORA in earlier generations, indicating that they prioritized the initial generation. Regarding the weights for GVP, GORA1 showed higher values than GORA2 and ORA, probably due to its higher initial values. We observed similar patterns under SI2 and SI3, with generally higher weights compared to SI1 (Supplementary Tables 1 and 2).

In Scenario 2, the comparison between GORA and ORA revealed trends similar to those in Scenario 1 (Supplementary Tables 3 - 5). However, the optimized weights were generally lower than those in Scenario 1 under identical selection strategies. Interestingly, the trend of prioritizing the initial generation—observed in GORAs for Scenario 1—was absent in Scenario 2. Conversely, ORA consistently placed greater importance on the final generation across all selection strategies. These results suggest that the optimized approaches flexibly changed their strategies depending on the target traits.

### 3.7 Optical flow for the gradients calculated by AD

Finally, to further validate the GORA strategies, we examined the global relationship between function values of *F* (***θ***) in equation (6) and gradients calculated by AD, i.e., *∂F/∂****θ***. We created visualizations similar to optical flow diagrams—a technique used in computer vision to represent object motion— (Horn and Schunck, 1981) for two parameters. We performed grid-based calculations of function values and gradients for two distinct breeding schemes based on two allocation parameters: (1) a scheme with one generation using both WBV and GVP, and (2) a scheme with two generations using only WBV for allocation across two scenarios under three selection strategies (Figure 5 for SI2 and Supplementary Figures 8 and 9 for SI1 and SI3). We defined the domains of logit-transformed weighting parameters ***ν***^(*τ*)^ as *ν*^(*τ*)^ ∈ [−3, 2] (*θ*^(*τ*)^ ∈ [−0.31, 3.02]) for WBV and *ν*^(*τ*)^ ∈ [−5, 0] (*θ*^(*τ*)^ ∈ [−0.47, 1.5]) for GVP. We then divided each domain into 50 areas and evaluated the function values and gradients for 51^2^ = 2, 601 parameter sets in total. In these visualizations, the background color indicates the magnitude of function values for each set of logit-transformed weighting parameters. The arrows, meanwhile, represent both the magnitude and direction of gradients calculated through AD.

**Fig. 5:**
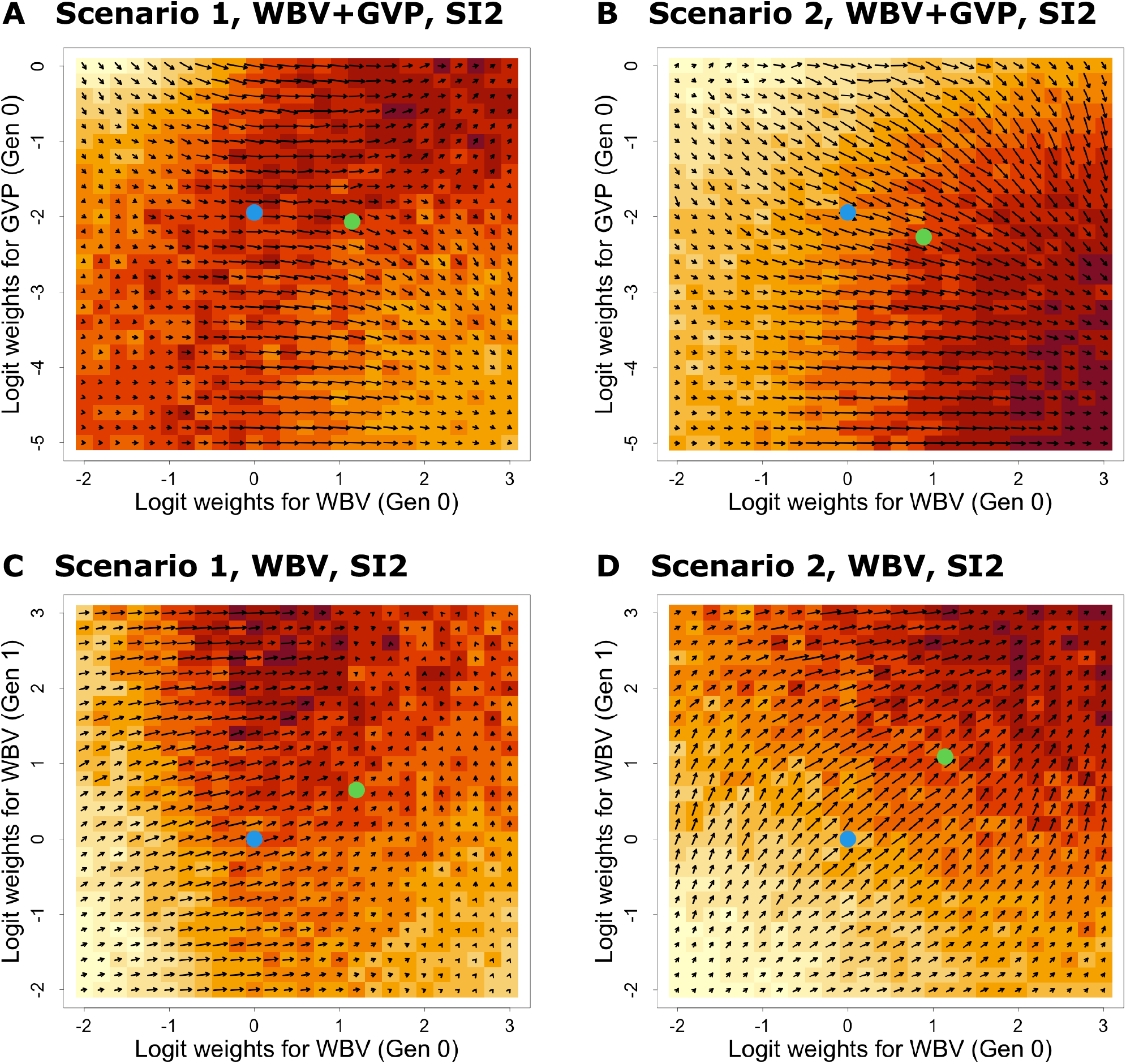
Optical flow for the gradients calculated by AD under SI2. The function values and gradients were evaluated by AD via grid search under SI2. The horizontal and vertical axes represent the logit-transformed weighting parameters ***ν***^(*τ*)^ in the allocation step. The background color indicates the magnitude of function values for each set of logit-transformed weighting parameters, whereas the arrows represent both the magnitude and direction of gradients calculated through AD. The blue and green points correspond to the initial and optimized values by the GORA2. (A), (B) The gradients were evaluated for the breeding scheme with only one generation considering both WBV and GVP as candidates of ***ω***_*k*_. The horizontal and vertical axes represent ***ν***^(*τ*)^ for WBV and GVP in generation 0, respectively. (C), (D) The gradients were evaluated for the breeding scheme with two generations considering only WBV as candidates of ***ω***_*k*_. The horizontal and vertical axes represent ***ν***^(*τ*)^ for WBV in generations 0 and 1, respectively. (A), (C) Scenario 1. (B), (D) Scenario 2.

Examining the results for the scheme with one generation using both WBV and GVP in Scenario 2, we observed that function values increased across all selection strategies as the weight for WBV grew larger and the weight for GVP diminished (Figure 5B and Supplementary Figures 8B and 9B). Also, the results for the scheme with two generations using only WBV in Scenario 2 showed that function values increased across all selection strategies as the weights for WBV in both generations got larger (Figure 5D and Supplementary Figures 8D and 9D). In both cases, the arrows representing gradient information pointed towards increasing function values, confirming that the gradient information was accurately calculated.

For the scheme with two generations using only WBV in Scenario 1, under SI3, we observed a trend similar to Scenario 2 (Supplementary Figure 9C). However, under SI1 and SI2, the highest function values occurred when the weight for WBV was low to moderate in generation 0 and high in generation 1 (Figure 5C and Supplementary Figure 8C). These findings suggest the optimal weighting parameters strongly depend on both QTL numbers and selection intensity the number of QTLs and selection intensity. Near the function’s maximum value, gradient arrows either pointed towards the peak or became negligible in length, ensuring that optimization would not stray from the optimal point. The scheme with one generation using both WBV and GVP in Scenario 1 revealed that optimal parameters varied significantly across selection strategies (Figure 5A and Supplementary Figures 8A and 9A). Notably, under SI2, a region of high function values seemed to extend along the diagonal line where x equals y (Figure 5A). In these cases, arrows primarily pointed rightward, sometimes failing to capture subtle changes in function values accurately.

The figures depict initial and optimized weighting parameters as blue and green points, respectively. In most cases, the SGD optimization search range was narrower than the grid search area in this subsection, suggesting it may not have reached the true optimal solution. Although this section assumes a shorter number of generations, these insights likely apply to breeding schemes with four generations as well, potentially explaining why GORA strategies showed slightly lower genetic gains than ORA.

## 4 Discussion

In this study, we calculated gradients from AD through DiffBreed, a PyTorch-based differentiable breeding simulator. We then leveraged these gradients to optimize progeny allocation in breeding schemes using gradient-based methods. We explored the relationship between function values and gradients across various parameter sets using grid search in breeding schemes spanning one or two generations (Figure 5 and Supplementary Figures 8 and 9). The optical flow diagrams showed that the arrows depicting gradients generally pointed towards regions of increasing function values. While capturing accurate gradients proved challenging for scenarios involving traits controlled by a small number of QTLs, our AD framework performed effectively for scenarios with traits governed by numerous QTLs. The number of QTLs influenced the continuity of genotypic values, which likely affected AD precision. In any case, based on these results, we can conclude that gradient estimation through AD achieved a satisfactory level of accuracy.

Having confirmed the validity of AD, we then applied the framework to the schemes with four generations. Monitoring function values across epochs in SGD revealed an increasing trend for most scenarios and selection intensities, with convergence typically occurring after about 100 epochs (Supplementary Figure 5). From these results, our gradient-based method with AD appeared to significantly reduce the number of breeding simulations required for optimization (100 *×* 200 = 20, 000) compared to the StoSOO-based optimization (50 *×* 20, 00 = 1000, 000) proposed by Hamazaki and Iwata (2024). However, it is important to note a key difference: our AD-based optimization used SGD, a local optimization method, while the previous study employed StoSOO (Valko et al., 2013) for global optimization. These approaches are fundamentally distinct, making direct comparisons of computational complexity inappropriate. Thus, the main focus of this study should be on how AD enabled more efficient and appropriate parameter exploration.

Using the optimized weighting parameters by SGD, we evaluated the performance of the proposed method based on the genetic gains (Figures 2 and 3 and Supplementary Figures 6 and 7). As a result, when the intensity in the selection stage was high, the optimized allocation strategies were unable to demonstrate clearly superior performance compared to non-optimized strategies. Although this result may seem discouraging initially, it is quite logical—the optimization of allocation strategies mainly works by enhancing selection intensity, so when the selection stage already employs high intensity, optimization provides only minimal additional benefit. While slightly inferior to StoSOO-based optimization, our gradient-based approach showed significantly better performance than non-optimized cases when selection intensity was low, confirming the effectiveness of our AD framework. Here, the reason why the gradient-based method yielded slightly less favorable results compared to the black-box-based method may be attributed to its limited range of exploration in the optimization process. As evident in the optical flow diagrams (Figure 5 and Supplementary Figures 8 and 9), even in breeding schemes spanning just one or two generations, the exploration range of SGD failed to encompass the entire global area, and consequently, SGD could not have reached the global optimum. One possible reason for the narrow search range of SGD is that we employed a scheduler, which might have prematurely halted the search process. Therefore, employing an alternative optimization method that can automatically adjust the learning rate, such as Adam (Kingma and Ba, 2014), without using a scheduler, could resolve this issue and potentially enhance the final genetic gains achieved through gradient-based optimization.

Next, we discussed the impact of different initial values on the performance of gradient-based methods. Analysis of genetic gains revealed that the gradient-based method with lower initial values for GVP consistently outperformed the approach with higher initial values (Figure 2 and Supplementary Figure 6). The results of genetic variance showed that higher initial GVP weights led to greater genetic diversity in breeding populations in all cases (Figure 4). These initial values significantly influenced the final optimized parameters (Table 1 and Supplementary Tables 1 - 5), likely explaining the observed variations in both genetic gains and diversity. These results indicate that initial values significantly impact the performance of gradient-based methods, even when applied to the area of breeding. Therefore, employing global optimization methods, such as Bayesian optimization (Brochu et al., 2010), to determine suitable initial values would enhance the effectiveness of our gradient-based method with AD.

Finally, the potential applications of AD extend far beyond simple gradient-based methods. In particular, black-box optimization methods that utilize derivative information—such as Bayesian optimization (Wu et al., 2017; Perrin and Leriche, 2024) or genetic algorithm (Sekhon and Mebane, 1998) with derivatives—show promise in accelerating optimization processes and enhancing their efficiency. By combining AD with derivative-oriented optimization methods, we can achieve greater genetic gains at lower computational costs, exceeding what our current gradient-based and black-box-based optimization frameworks can deliver. Moreover, the gradient information obtained through AD can be incorporated into deep learning techniques, such as deep neural networks (Schmidhuber, 2015). By integrating deep neural network structures into the optimization of breeding schemes using AD, we could potentially create a more flexible optimization framework that does not rely on human-proposed goodness metrics and can handle a larger number of parameters. Thus, the DiffBreed framework developed in this study is expected to serve as a foundation for more sophisticated optimization methods in the future. Furthermore, our AD framework could be extended beyond breeding applications to other fields, including simulations in population genetics, as suggested in Frank (2022). By establishing the AD of breeding schemes and demonstrating its potential through gradient-based optimization, this study lays a crucial foundation for revolutionizing future decision-making in breeding strategies.

## Author contributions statement

K.H., H.I., and K.T. conceived and designed the study. K.H. implemented and developed the proposed framework. K.H. and K.T. drafted the manuscript. K.T. supervised the study. All authors have read and approved the final manuscript.

## Acknowledgments

This work was supported by JST, ACT-X Grant Number JPMJAX23CL, Japan. The computations in this work were performed in the supercomputer centers at RAIDEN. We would like to thank Dr. Masato Sumita for his assistance in setting up the computational environment such as RAIDEN.

## References

A. Allier, C. Lehermeier, A. Charcosset, L. Moreau, and S. Teyssédre. Improving short and long-term genetic gain by accounting for within-family variance in optimal cross-selection. Front. Genet., 10:1006, Oct. 2019a.

A. Allier, L. Moreau, A. Charcosset, S. Teyssédre, and C. Lehermeier. Usefulness criterion and post-selection parental contributions in multi-parental crosses: Application to polygenic trait introgression. G3, 9(5):1469–1479, May 2019b.

M. AlQuraishi and P. K. Sorger. Differentiable biology: using deep learning for biophysics-based and data-driven modeling of molecular mechanisms. Nat. Methods, 18(10):1169–1180, Oct. 2021.

A. G. Baydin, B. A. Pearlmutter, A. A. Radul, and J. M. Siskind. Automatic differentiation in machine learning: A survey. J. Mach. Learn. Res., 18(153):1–43, 2018.

J. Bradbury, R. Frostig, P. Hawkins, M. J. Johnson, C. Leary, D. Maclaurin, G. Necula, A. Paszke, J. VanderPlas, S. Wanderman-Milne, and Q. Zhang. JAX: Autograd and XLA, Nov. 2021.

E. Brochu, V. M. Cora, and N. de Freitas. A tutorial on bayesian optimization of expensive cost functions, with application to active user modeling and hierarchical reinforcement learning. arXiv [cs.LG], Dec. 2010.

A. Callejo, S. H. K. Narayanan, J. García de Jalón, and B. Norris. Performance of automatic differentiation tools in the dynamic simulation of multibody systems. Adv. Eng. Softw., 73:35–44, July 2014.

J. L. Capper. Replacing rose-tinted spectacles with a high-powered microscope: The historical versus modern carbon footprint of animal agriculture. Anim. Front, 1(1):26–32, July 2011.

J. Diot. breedSimulatR: R Breeding Simulator, 2025. https://github.com/ut-biomet/breedSimulatR, https://ut-biomet.github.io/breedSimulatR/.

J. Diot and H. Iwata. Bayesian optimisation for breeding schemes. Front. Plant Sci., 13:1050198, 2022.

S. A. Frank. Automatic differentiation and the optimization of differential equation models in biology. Front. Ecol. Evol., 10: 1010278, Nov. 2022.

R. C. Gaynor, G. Gorjanc, and J. M. Hickey. AlphaSimR: an R package for breeding program simulations. G3 (Bethesda), 11 (2):jkaa017, Feb. 2021.

M. Goddard. Genomic selection: prediction of accuracy and maximisation of long term response. Genetica, 136(2):245–257, June 2009.

A. Griewank and A. Walther. Evaluating derivatives: principles and techniques of algorithmic differentiation: Principles and techniques of algorithmic differentiation, second edition. Cambridge University Press, Cambridge, TAS, Australia, 2 edition, 2008.

K. Hamazaki and H. Iwata. AI-assisted selection of mating pairs through simulation-based optimized progeny allocation strategies in plant breeding. Front. Plant Sci., 15, 2024.

B. K. P. Horn and B. G. Schunck. Determining optical flow. Artif. Intell., 17(1-3):185–203, Aug. 1981.

S. R. Hunter and B. McClosky. Maximizing quantitative traits in the mating design problem via simulation-based pareto estimation. IIE Trans., 48(6):565–578, June 2016.

E. Jang, S. Gu, and B. Poole. Categorical reparameterization with gumbel-softmax. arXiv [stat.ML], Nov. 2016.

J.-L. Jannink. Dynamics of long-term genomic selection. Genet. Sel. Evol., 42:35, Aug. 2010.

J.-L. Jannink, R. Astudillo, and P. Frazier. Insight into a two-part plant breeding scheme through bayesian optimization of budget allocations. Crop Sci., Oct. 2023.

D. P. Kingma and J. Ba. Adam: A method for stochastic optimization. arXiv [cs.LG], Dec. 2014.

C. Lauvernet, L. Hascoët, F.-X. Le Dimet, and F. Baret. Using automatic differentiation to study the sensitivity of a crop model. In Lecture Notes in Computational Science and Engineering, Lecture notes in computational science and engineering, pages 59–69. Springer Berlin Heidelberg, Berlin, Heidelberg, 2012.

C. Lehermeier, S. Teyssédre, and C.-C. Schön. Genetic gain increases by applying the usefulness criterion with improved variance prediction in selection of crosses. Genetics, 207(4): 1651–1661, Dec. 2017.

I. A. Luchnikov, M. E. Krechetov, and S. N. Filippov. Riemannian geometry and automatic differentiation for optimization problems of quantum physics and quantum technologies. New J. Phys., 23(7):073006, July 2021.

C. J. Maddison, A. Mnih, and Y. W. Teh. The concrete distribution: A continuous relaxation of discrete random variables. In International Conference on Learning Representations, 2017.

Martín Abadi, Ashish Agarwal, Paul Barham, Eugene Brevdo, Zhifeng Chen, Craig Citro, Greg S. Corrado, Andy Davis, Jeffrey Dean, Matthieu Devin, Sanjay Ghemawat, Ian Goodfellow, Andrew Harp, Geoffrey Irving, Michael Isard, Y. Jia, Rafal Jozefowicz, Lukasz Kaiser, Manjunath Kudlur, Josh Levenberg, Dandelion Mané, Rajat Monga, Sherry Moore, Derek Murray, Chris Olah, Mike Schuster, Jonathon Shlens, Benoit Steiner, Ilya Sutskever, Kunal Talwar, Paul Tucker, Vincent Vanhoucke, Vijay Vasudevan, Fernanda Viégas, Oriol Vinyals, Pete Warden, Martin Wattenberg, Martin Wicke, Yuan Yu, and Xiaoqiang Zheng. TensorFlow: Large-scale machine learning on heterogeneous systems, 2015.

T. H. Meuwissen. Maximizing the response of selection with a predefined rate of inbreeding. J. Anim. Sci., 75(4):934–940, Apr. 1997.

S. Moeinizade, G. Hu, L. Wang, and P. S. Schnable. Optimizing selection and mating in genomic selection with a look-ahead approach: An operations research framework. G3, 9(7):2123– 2133, July 2019.

S. Moeinizade, G. Hu, and L. Wang. A reinforcement learning approach to resource allocation in genomic selection. Intelligent Systems with Applications, 14:200076, May 2022.

A. Paszke, S. Gross, S. Chintala, G. Chanan, E. Yang, Z. DeVito, Z. Lin, A. Desmaison, L. Antiga, and A. Lerer. Automatic differentiation in pytorch. In NIPS 2017 Workshop on Autodiff, 2017. URL https://openreview.net/forum?id=BJJsrmfCZ.

A. Paszke, S. Gross, F. Massa, A. Lerer, J. Bradbury, G. Chanan, T. Killeen, Z. Lin, N. Gimelshein, L. Antiga, A. Desmaison, A. Köpf, E. Yang, Z. DeVito, M. Raison, A. Tejani, S. Chilamkurthy, B. Steiner, L. Fang, J. Bai, and S. Chintala. PyTorch: An imperative style, high-performance deep learning library. Adv. Neural Inf. Process. Syst., abs/1912.01703, Dec. 2019.

G. Perrin and R. Leriche. Bayesian optimization with derivatives acceleration. Transactions on Machine Learning Research, Apr. 2024.

R Core Team. R: A language and environment for statistical computing, 2024.

L. B. Rall. Automatic Differentiation: Techniques and Applications. Lecture notes in computer science. Springer, Berlin, Germany, 1981 edition, Aug. 1981.

H. Robbins and S. Monro. A stochastic approximation method. aoms, 22(3):400–407, Sept. 1951.

K. Sakurai, K. Hamazaki, M. Inamori, A. Kaga, and H. Iwata. Cross potential selection: A proposal for optimizing crossing combinations in recurrent selection using the usefulness criterion of future inbred lines. G3 (Bethesda), page jkae224, Sept. 2024.

J. Schmidhuber. Deep learning in neural networks: an overview. Neural Netw., 61:85–117, Jan. 2015.

J. S. Sekhon and W. R. Mebane, Jr. Genetic optimization using derivatives. Polit. Anal., 7:187–210, Jan. 1998.

R. Z. Shrote and A. M. Thompson. PyBrOpS: a python package for breeding program simulation and optimization for multi-objective breeding. G3 (Bethesda), 14(10):jkae199, Oct. 2024.

T. Tamayo-Mendoza, C. Kreisbeck, R. Lindh, and A. Aspuru Guzik. Automatic differentiation in quantum chemistry with applications to fully variational hartree-fock. ACS Cent. Sci., 4 (5):559–566, May 2018.

M. Valko, A. Carpentier, and R. Munos. Stochastic simultaneous optimistic optimization. In S. Dasgupta and D. McAllester, editors, Proceedings of the 30th International Conference on Machine Learning, volume 28 of Proceedings of Machine Learning Research, pages 19–27, Atlanta, Georgia, USA, 2013. PMLR.

G. Van Rossum and F. L. Drake. Python 3 Reference Manual. CreateSpace, Scotts Valley, CA, 2009.

J. Wu, M. Poloczek, A. G. Wilson, and P. I. Frazier. Bayesian optimization with gradients. arXiv [stat.ML], Mar. 2017.

N. Yoshikawa and M. Sumita. Automatic differentiation for the direct minimization approach to the hartree-fock method. J. Phys. Chem. A, 126(45):8487–8493, Nov. 2022.

