## Supplementary Data 1 for "DiffBreed: Automatic differentiation enables efficient gradient-based optimization of breeding strategies"

Short title: DiffBreed: gradient-based optimization of breeding

Kosuke Hamazaki[1*], Hiroyoshi Iwata[2], and Koji Tsuda[1,3*]

[1] Molecular Informatics Team, RIKEN Center for Advanced Intelligence Project (AIP), RIKEN, 178-4-4 Wakashiba, Kashiwa, Chiba 277-0871, Japan.

[2] Laboratory of Biometry and Bioinformatics, Department of Agricultural and Environmental Biology, Graduate School of Agricultural and Life Sciences, The University of Tokyo, 1-1-1 Yayoi, Bunkyo-Ku, Tokyo 113-8657, Japan.

[3] Department of Computational Biology and Medical Sciences, Graduate School of Frontier Sciences, The University of Tokyo, 5-1-5 Kashiwa-no-ha, Kashiwa, Chiba 277-8561, Japan.

* Corresponding author

 (KH), (KT)

### Supplementary Sections

#### S1 Details of simulating the initial breeding population and QTLs

##### S1.1 Details of simulating genome structure in the initial breeding population

The marker genotype of the initial population was simulated similarly to Hamazaki and Iwata (2024) . As in their study, we assumed true QTL positions and effects were known for simplicity. In the simulation, only genome-wide QTLs for founder haplotypes were generated using the coalescent simulator GENOME (Liang *et al.*, 2007). This study assumed a virtual diploid crop with ten chromosomes ($2n = 20$) and $m^{(\mathrm{Chr})}$ QTLs on each chromosome. Here, we prepared two different scenarios concerning the number of QTLs: $m^{(\mathrm{Chr})} = 3$ (Scenario 1) or $m^{(\mathrm{Chr})} = 50$ (Scenario 2).

First, 4,000 founder haplotypes for each chromosome were independently generated using GENOME with the following parameters: "-pop 1 4000 -N 4000 -c 1 -pieces 20000 -rec 0.0001 -s 40 -maf 0.01 -tree 0 -mut 0.00000001" (Scenario 1) or "-pop 1 4000 -N 4000 -c 1 -pieces 20000 -rec 0.0001 -s 416 -maf 0.01 -tree 0 -mut 0.00000001" (Scenario 2). Among the $8\left(m^{(\mathrm{QTL})} + 2\right)$ loci, $m^{(\mathrm{Chr})}$ loci whose minor allele frequency (MAF) was equal to or larger than 0.01 were randomly selected as QTLs. Next, 2,000 founders were generated by sampling two haplotypes from all founder haplotypes with replacement. These founders were randomly mated for one generation to simulate a historical population of the 2,000 genotypes. These procedures for simulating the historical population are similar to those in the R package "BreedingSchemeLanguage" by Yabe *et al.* (2017) .

Then, the base breeding population was simulated from this historical population in a similar manner as previous research (Müller *et al.*, 2017, 2018). Concretely, the historical population underwent random mating for 300 generations, followed by a population bottleneck that randomly selected 200 individuals and conducted random mating for 15 generations while maintaining a population size of 2,000. This bottleneck was used to create extensive linkage disequilibrium (LD), often observed in elite plant breeding populations (Van Inghelandt *et al.*, 2011). The population was then subject to random mating for three more generations to remove close family relationships, resulting in pre-breeding materials.

From these simulated pre-breeding materials, $N = 250$ genotypes were selected for the initial population of a breeding scheme so that these 250 genotypes represented the simulated genetic resources in terms of genetic diversity. This selection was conducted in the completely same manner as Hamazaki and Iwata (2024) , i.e., using the $k$-medoids method on the marker genotypes of the pre-breeding material using the "pam" function of the R package "cluster" version 2.1.2 (Maechler *et al.*, 2021). After clustering the pre-breeding materials into $250$ groups, we selected the medoids as the representative genotypes from each group for the initial breeding population. This base breeding population was regarded as generation $t = 0$ in a breeding scheme.

#### S1.2 Details of simulating QTLs and genotypic values

We assumed quantitative traits with a simple genetic architecture as target traits in this study. First, as described in the previous subsection, $m^{(\mathrm{Chr})} = 3$ or $m^{(\mathrm{Chr})} = 50$

QTLs were assumed to have some effects on phenotypes (i.e., a total of $m = 30$ or $m = 500$ QTLs). QTL effects were then sampled from the normal distribution, as shown in equation (S1).

$$\mathbf{a} \sim \mathrm{MVN}\left(\mathbf{0}, \frac{1}{m}\mathbf{I}_m\right), \tag{S1}$$

where $\mathbf{a} \in \Re^m$ is a vector of QTL effects and $\mathbf{I}_m \in \Re^{m \times m}$ is an identity matrix. Here, all QTL effects were assumed to be additive for simplicity. Then, $\boldsymbol{\alpha} \in \Re^M$ in the main manuscript was created by duplicating $\mathbf{a}$, i.e., $\boldsymbol{\alpha} = [\mathbf{a}^\top, \mathbf{a}^\top]^\top$ where $M = 2m$. The true additive genotypic value $u_i$ for the parent panel were then simulated using the following equation (S2).

$$u_i = \boldsymbol{\alpha}^\top \mathbf{w}_i, \tag{S2}$$

where $\mathbf{w}_i \in \Re^M$ is a vector of allele scores of an individual $i$ formed by vertically stacking two haplotypes. The additive genotypic values in equaltion (S2) was utilized to evaluate the population quality $F$ in equation (1).

### S2 Details of breeding schemes

Here, we describe more details of the breeding scheme assumed in this study. The breeding scheme consists of the following steps:

1. Select parent candidates (Supplementary Figure 4 (1))

   Based on the true marker (QTL) effects, we first computed WBVs for individuals in the current generation $t$ ($t \in \{0, \dots, T-1\}$) to select parent candidates as in the following equation (S3) (Jannink, 2010).

$$\varphi_i = \sum_{j=1}^{M} \alpha_j p_j^{-\frac{1}{2}} w_{ij} , \quad \text{(S3)}$$

where $\varphi_i$ is a WBV of an individual $i$ and the other terms are defined in equation (8). WBV has been expected to maintain genetic diversity by emphasizing rare allele effects and thus avoid the saturation of genetic gains in mid- or long-term breeding programs compared to BV (Goddard, 2009; Jannink, 2010).

Then, based on the order of increasing WBV, the $n$ genotypes were selected as parent candidates for mating. In this study, we implemented the three selection methods regarding selection intensity, as described in the main manuscript.

2. Determine mating pairs for the next generation (Supplementary Figure 4 (2))

The selected $n$ parent candidates were then subjected to diallel crossing, including selfing, to determine the mating pairs for the next generation. This process yielded $k = \frac{n(n+1)}{2}$ total crossing pairs.

3. Allocate progenies to each mating pair (Supplementary Figure 4 (3))

Three strategies were employed to allocate progenies to each mating pair determined in Step 2, after which a crossing table was created. Here, as described in the main manuscript, to account for crossing costs, we implemented a constraint requiring that progeny allocations for crosses be made in minimum units of $L = 5$ individuals. Additionally, all strategies produced $N$ progenies in total for the next generation. In this study, we assumed $N$ remained constant over generations, i.e., $N = 250$.

a. The gradient-based optimized resource allocation method (GORA)

The first strategy was the gradient-based optimized allocation method proposed in this study. We enhanced the progeny allocation proposed in Hamazaki and Iwata (2024) by incorporating the Gumbel-Softmax function (Jang *et al.*, 2017;

Maddison *et al.*, 2017), extending its applicability to the AD framework.

b. The black-box-based optimized resource allocation method (ORA)

The second strategy was the optimized allocation method proposed in Hamazaki and Iwata (2024). This method applied the softmax function to the weighted sum of the selection criteria computed in Step 1. Then, the weights for the criteria were optimized by the black-box optimization method through repeated breeding simulations.

c. Equal resource allocation method (EQ)

We also implemented an equal allocation method that allocated $L$ progenies each to $\frac{N}{L}$ pairs, which were randomly selected from the $k$ candidate pairs.

4. Generate progenies for the next generation (Supplementary Figure 4 (4))

Following the crossing table created in Step 3, two gametes were generated for each mating pair based on the method described in Section 2.2, accounting for marker recombination. Recombination rates between markers were calculated using the Kosambi map function (Kosambi, 1943; Zhao and Speed, 1996) based on the linkage map generated by GENOME. These two gametes were then combined to form one new progeny, resulting in a new population which consisted (with generation $t+1$) of $N$ progenies.

5. Repeat the parent selection and mating process

Steps 1-4 were repeated until the population reached the final generation, $t = T$. This process was carried out for $T - 1$ generations, where $T$ represents the total number of generations in the breeding program.

Finally, we evaluated the results for each strategy using the true additive genotypic values of the final population with generation $T$, $\mathbf{u}^{(T)}$. In this study, similar to Hamazaki and Iwata (2024), we set the final generation as $T = 4$, assuming a scenario where rapid genetic improvement was needed in small-scale breeding schemes. Here, to evaluate the population's maximum for the breeding strategy, we computed $F^{(t)}$, the empirical mean of $\mathbf{u}^{(t)}$ for the top $K$ genotypes as in equation (1). These genotypes were selected based on their $\mathbf{u}^{(t)}$ values in descending order. We then scaled $F^{(t)}$ based on the initial population's mean and standard deviation, $\bar{F}^{(0)}$ and $\sigma_u^{(0)}$, resulting $\tilde{F}^{(t)} = \left(F^{(t)} - \bar{F}^{(0)}\right)/\sigma_u^{(0)}$. We chose $K = 5$ to represent the top individuals for new variety development while mitigating stochastic variation between simulations. For each parent panel simulation dataset, we ran 10,000 different breeding schemes per strategy and calculated the empirical mean of $\tilde{F}^{(t)}$ from these results. We repeated this simulation and evaluation process for 10 replicates of the phenotype simulation, each with different QTL positions and effects.

### S3 Gradient-based optimization of allocation strategies

In this study, we leveraged gradient information obtained through AD of breeding simulations implemented in PyTorch for gradient-based optimization methods (Supplementary Figure S1). Among various gradient techniques, we employed Stochastic Gradient Descent (SGD) (Robbins and Monro, 1951) with an initial learning rate of 0.15 to optimize the allocation strategy. To ensure proper control of the learning rate in the latter stages of optimization, we applied a learning rate scheduler. Using PyTorch's torch.optim.lr_scheduler.StepLR function (Paszke *et al.*, 2019), we decreased the learning rate by a factor of 0.8 every 10 epochs. As described in Section 3.1, we conducted 100 breeding simulations for each parameter set evaluation to obtain gradient information while dealing with the randomness caused by progeny generation. Here, we defined the domain of definition using the inverse logit transformation as shown in equation (S4).

$$\boldsymbol{\theta}^{(\tau)} = (\theta_{\max} - \theta_{\min}) \times \frac{\exp\left(\boldsymbol{\nu}^{(\tau)}\right)}{1 + \exp\left(\boldsymbol{\nu}^{(\tau)}\right)} + \theta_{\min}, \tag{S4}$$

where $\theta_{\max} = 3.5$ and $\theta_{\min} = -0.5$ are the minimum and maximum values of $\boldsymbol{\theta}^{(\tau)}$, respectively, and $\boldsymbol{\nu}^{(\tau)} \in (-\infty, \infty)$ is logit-transformed weighting parameter, serving as the actual input to the function and the target of optimization. Then, we set 1.5 as the initial parameter for each element of $\left\{\boldsymbol{\theta}^{(\tau)}\right\}_{\tau=0,\ldots,T-1}$ (GORA1). To assess the impact of initial values on our results, we included a strategy where the initial weight for GVP was set to 0 (GORA2). We then repeated these function evaluations 200 times, updating parameters via SGD. The final parameter set after these 200 epochs was considered the optimal set of parameters. These GORA approaches were also evaluated under the constraint with regard to the minimum number of allocated progenies to each pair to compare GORAs on an equal footing with the other strategies.

### Supplementary Figures

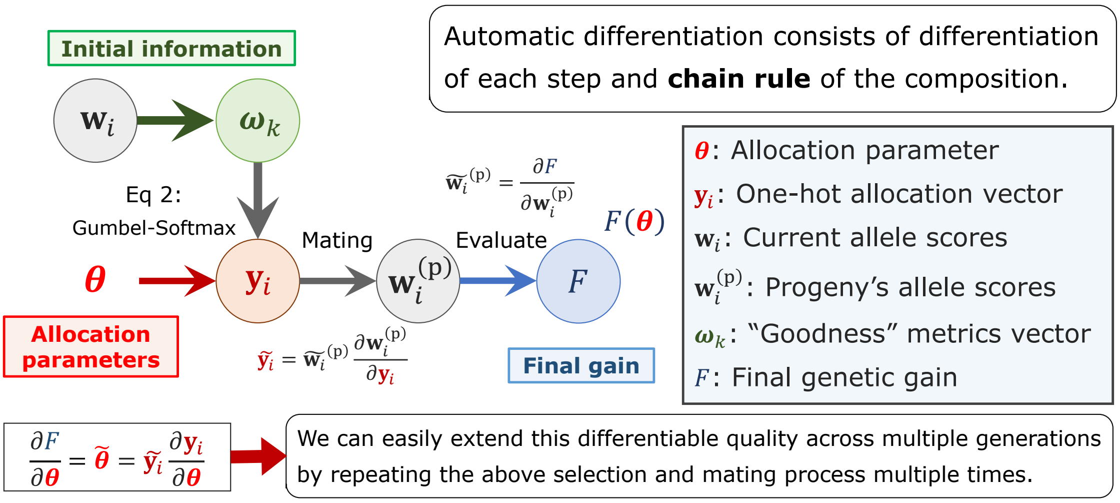


**Supplementary Figure 1. Example of AD of the genetic gain in next generation with respect to the allocation parameters.**

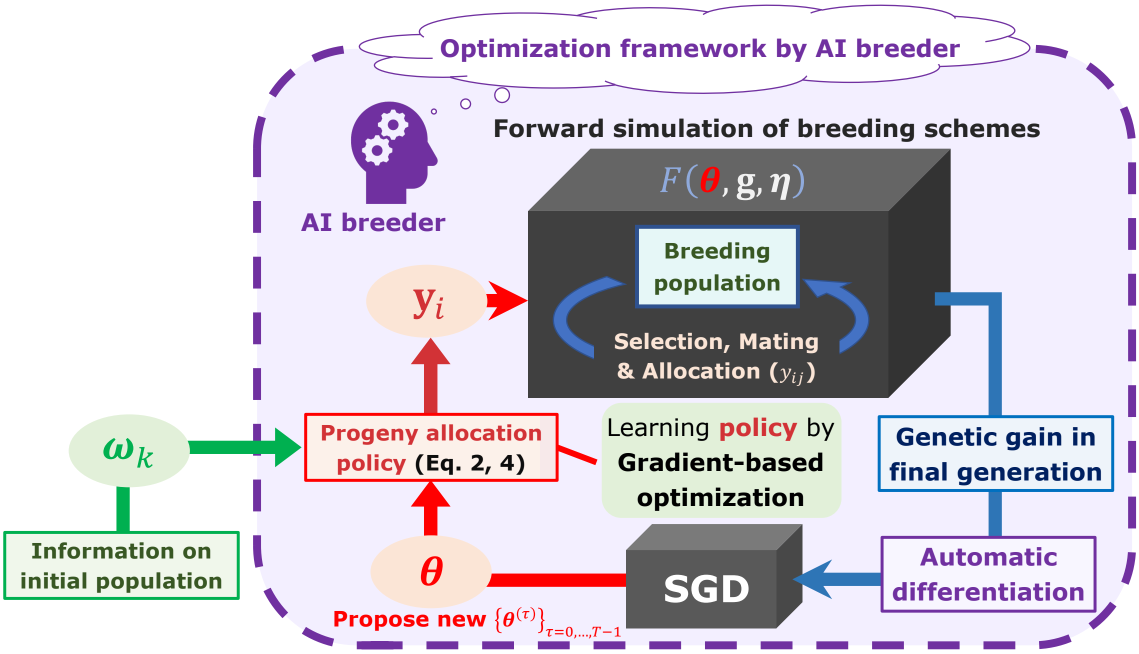


**Supplementary Figure 2. The optimization framework of allocation strategy in the breeding scheme.** In our framework, the breeders can obtain a set of parameters representing the optimal allocation strategy, $\{\widehat{\boldsymbol{\theta}}^{(\tau)}\}_{\tau=0,\ldots,T-1}$, by providing the AI breeder with information on an initial breeding population, as in (Hamazaki and Iwata, 2024). Here, a breeding simulator is regarded as a function whose input is a set of allocation parameters, and whose output is the final genetic gain. After computing the function derivative by using the AD technique with PyTorch, the allocation parameters are optimized by the gradient-based approach such as SGD.

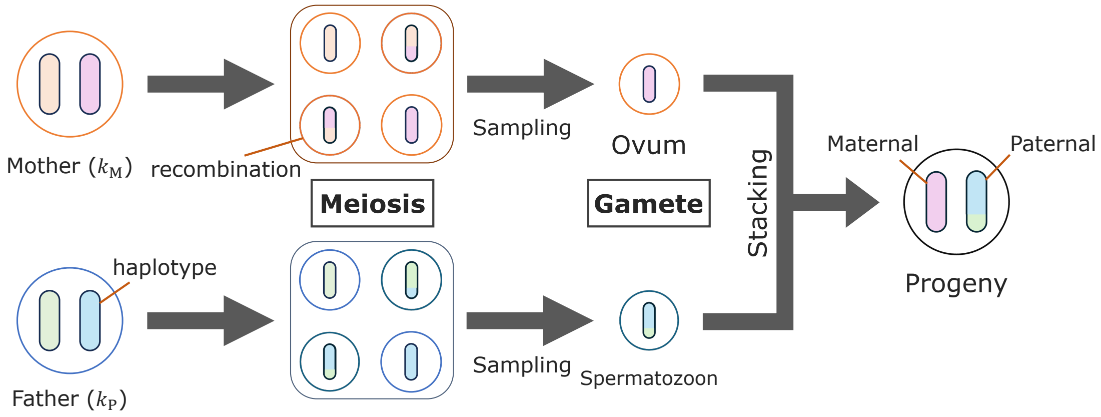


**Supplementary Figure 3. Image of generating progeny via meiosis.**

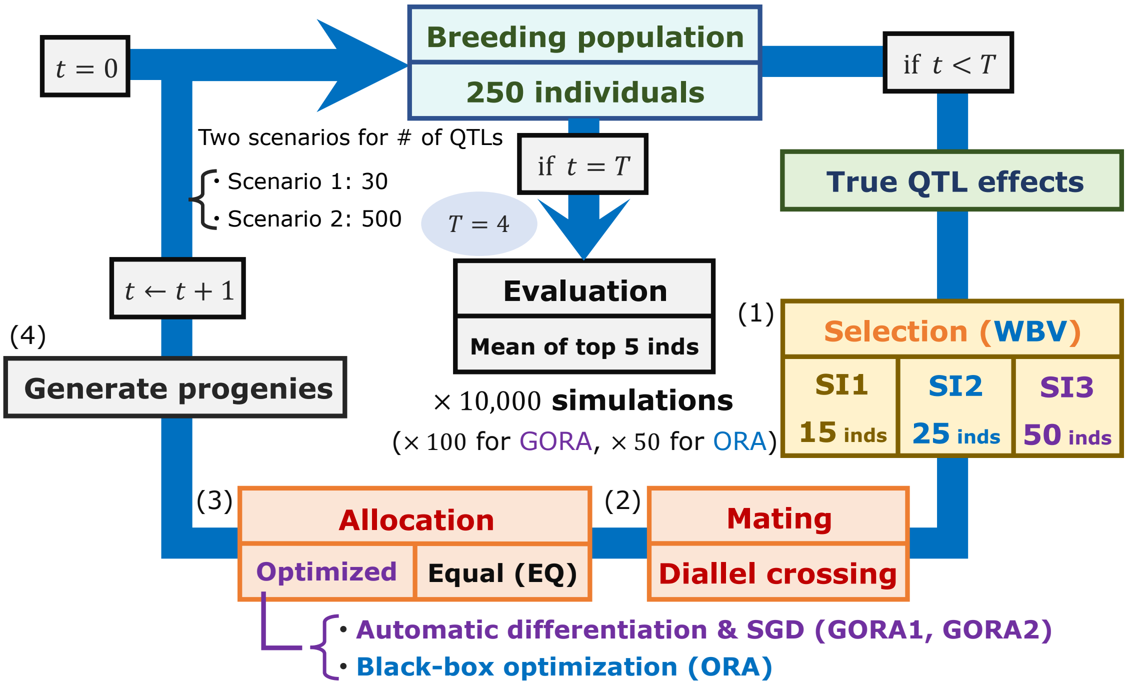


**Supplementary Figure 4. Breeding schemes assumed in this study.** Similar to Hamazaki and Iwata (2024), we assumed small-scale plant breeding programs with recurrent selection. In a scheme, the selection, mating, and allocation steps were repeated until the final generation. Then, the gradient-based optimized allocation (GORA) strategy was compared with the black-box-based optimized allocation (ORA) and non-optimized equal allocation (EQ) strategies. Here, we prepared for the two scenarios with different numbers of QTLs and three selection strategies with different selection intensities.

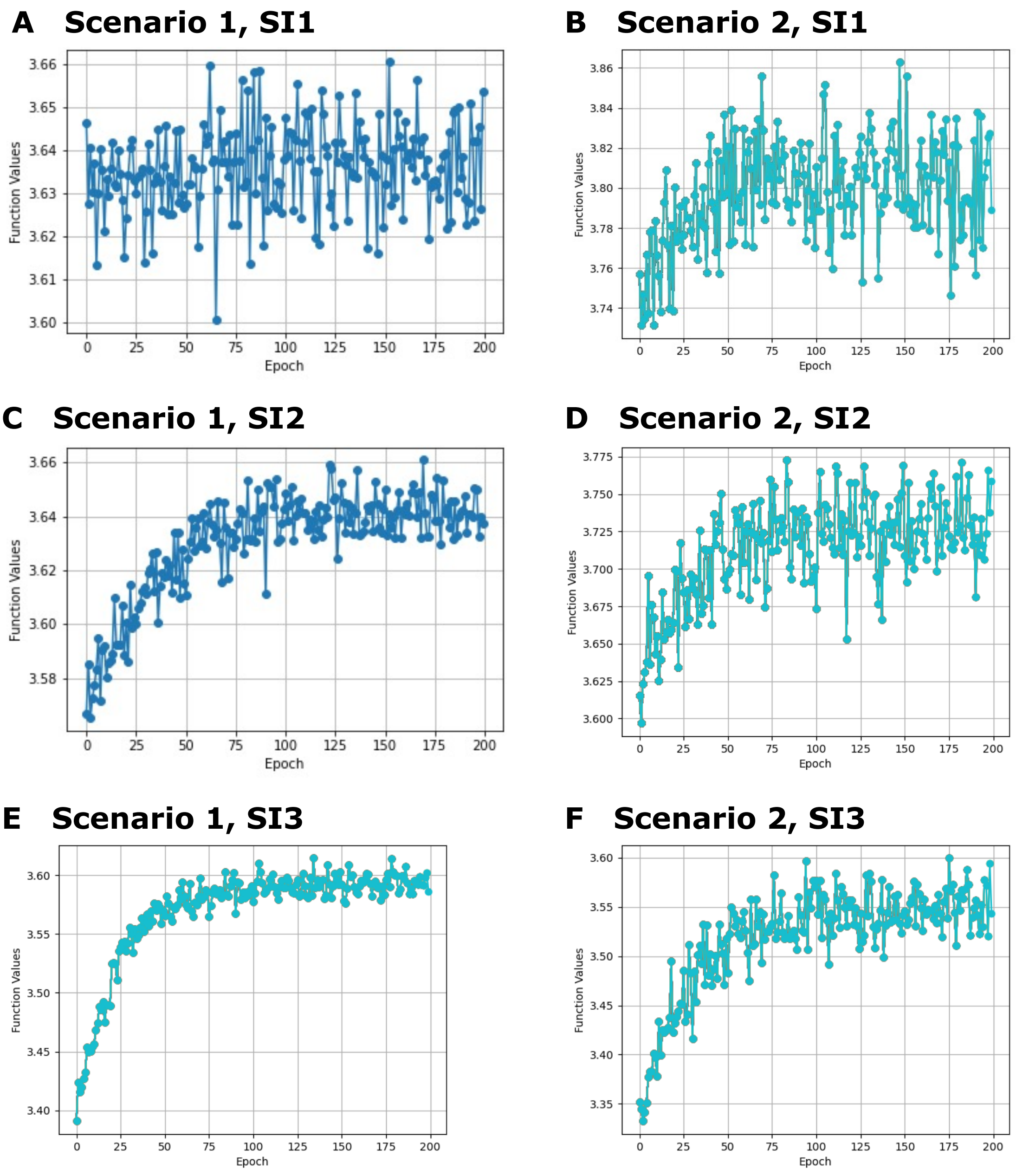


**Supplementary Figure 5. Convergence conditions of the SGD algorithm.** Change in the function values with different epochs. The horizontal and vertical axes represent the number of epochs of the SGD algorithm and the function values evaluated with 100 breeding schemes, respectively. (A), (C), (E) Scenario 1. (B), (D), (F) Scenario 2. (A), (B) SI1. (C), (D) SI2. (E), (F) SI3.

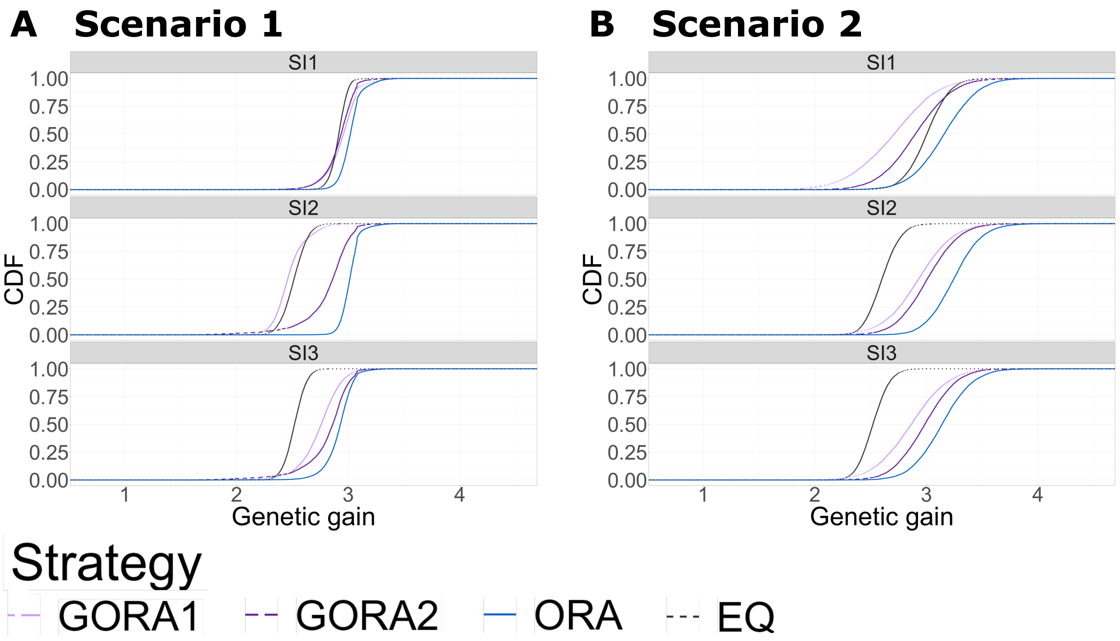


**Supplementary Figure 6. Genetic gains across different simulation repetitions.** The horizontal axis shows the final genetic gain of an individual, whereas the vertical axis represents the percentile of the simulation repetitions. We compared the four allocation strategies under the different selection intensities (SI1 – SI3). The abbreviations of the allocation strategies are the same as those of Figure 1. (A) Scenario 1. (B) Scenario 2.

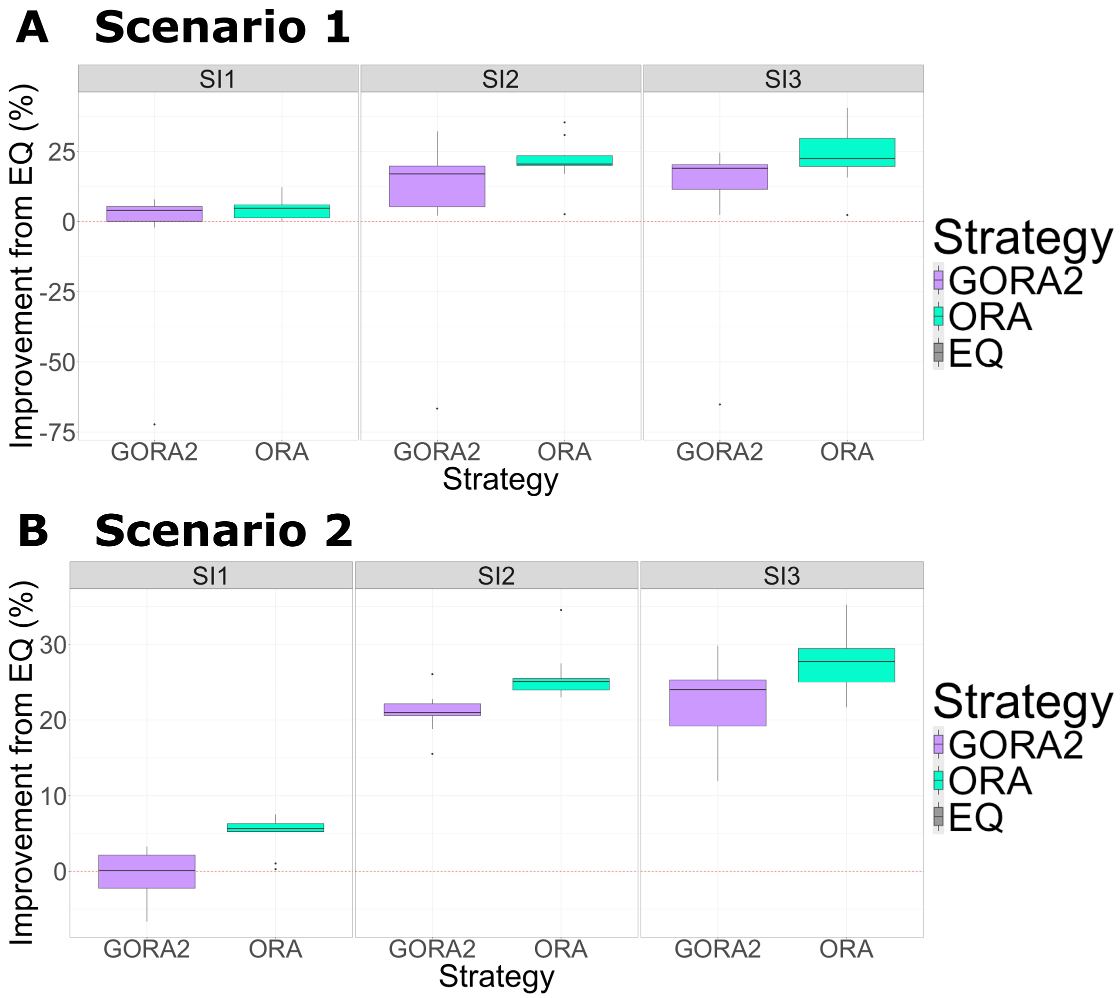


**Supplementary Figure 7. Improvement of the final genetic gains compared to the equal allocation strategy using ten replications for the phenotype simulation under different selection intensities.** Boxplots of the improvement of the final genetic gains compared to the equal allocation strategy using ten replications for the phenotype simulation. The horizontal and vertical axes represent the different allocation strategies and the improvement rate of the final genetic gains, respectively. We compared the two allocation strategies under the different selection intensities (SI1 – SI3): GORA2 (light purple) and ORA (light blue), with details given in Figure 2. The weighting parameters were optimized based on 100 and 5,000 function evaluations for GORA2 and ORA, respectively. (A) Scenario 1. (B) Scenario 2.

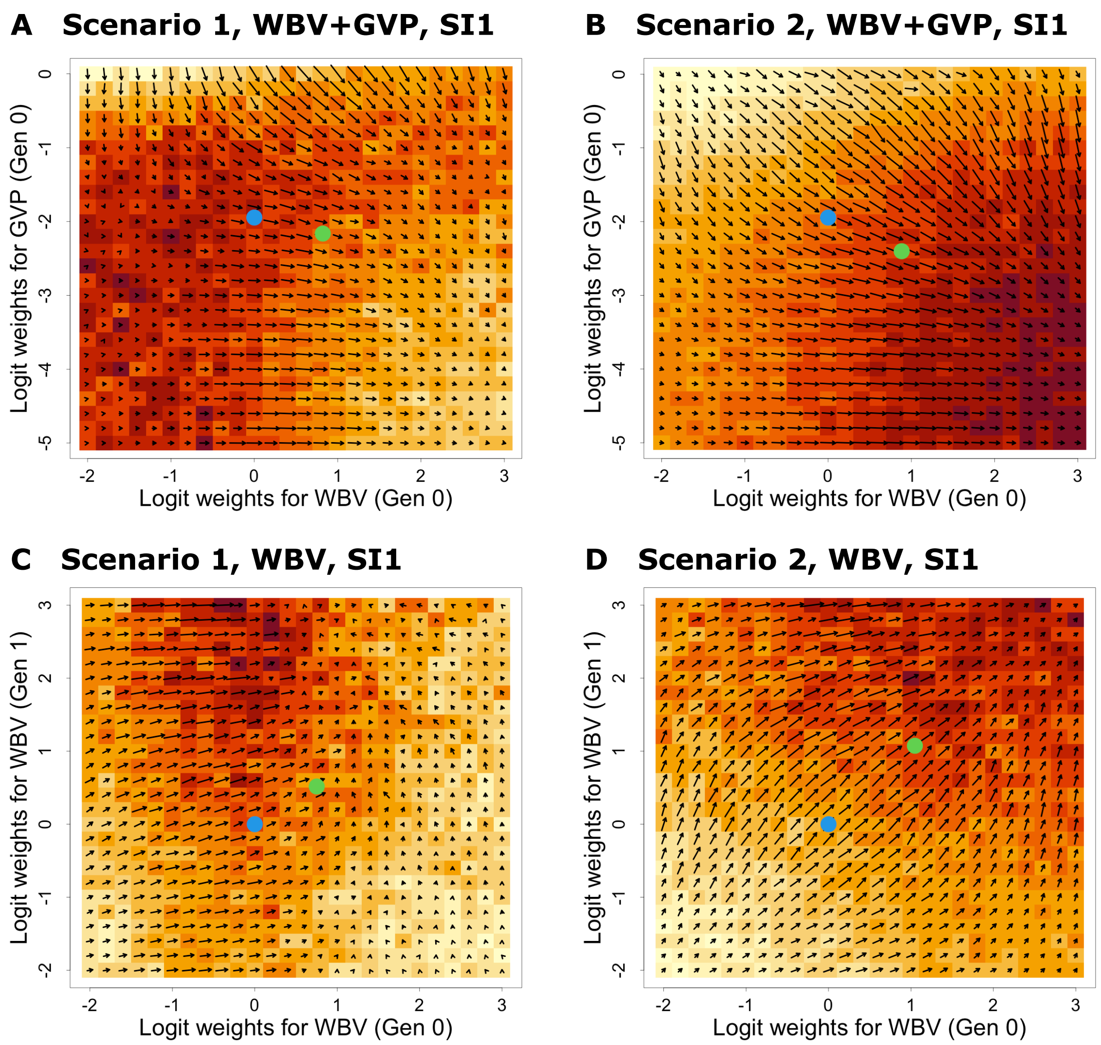


**Supplementary Figure 8. Optical flow for the gradients calculated by AD under SI1.** The function values and gradients were evaluated by AD via grid search under SI1. The horizontal and vertical axes represent the logit-transformed weighting parameters $\boldsymbol{\nu}^{(\tau)}$ in the allocation step. The background color represents the size of the function values for each set of logit-transformed weighting parameters, whereas the arrows reflect the sizes and directions of the gradients evaluated by AD. The blue and green points correspond to the initial and optimized values by the GORA2 approach. (A), (B) The gradients were evaluated for the breeding scheme with only one generation considering both WBV and GVP as candidates of $\boldsymbol{\omega}$. The horizontal and vertical axes represent the logit-transformed weighting parameters for WBV and GVP in generation 0, respectively. (C), (D) The gradients were evaluated for the breeding scheme with two generations considering only WBV as candidates of $\boldsymbol{\omega}$. The horizontal and vertical axes represent the logit-transformed weighting parameters for WBV in generations 0 and 1, respectively. (A), (C) Scenario 1. (B), (D) Scenario 2.

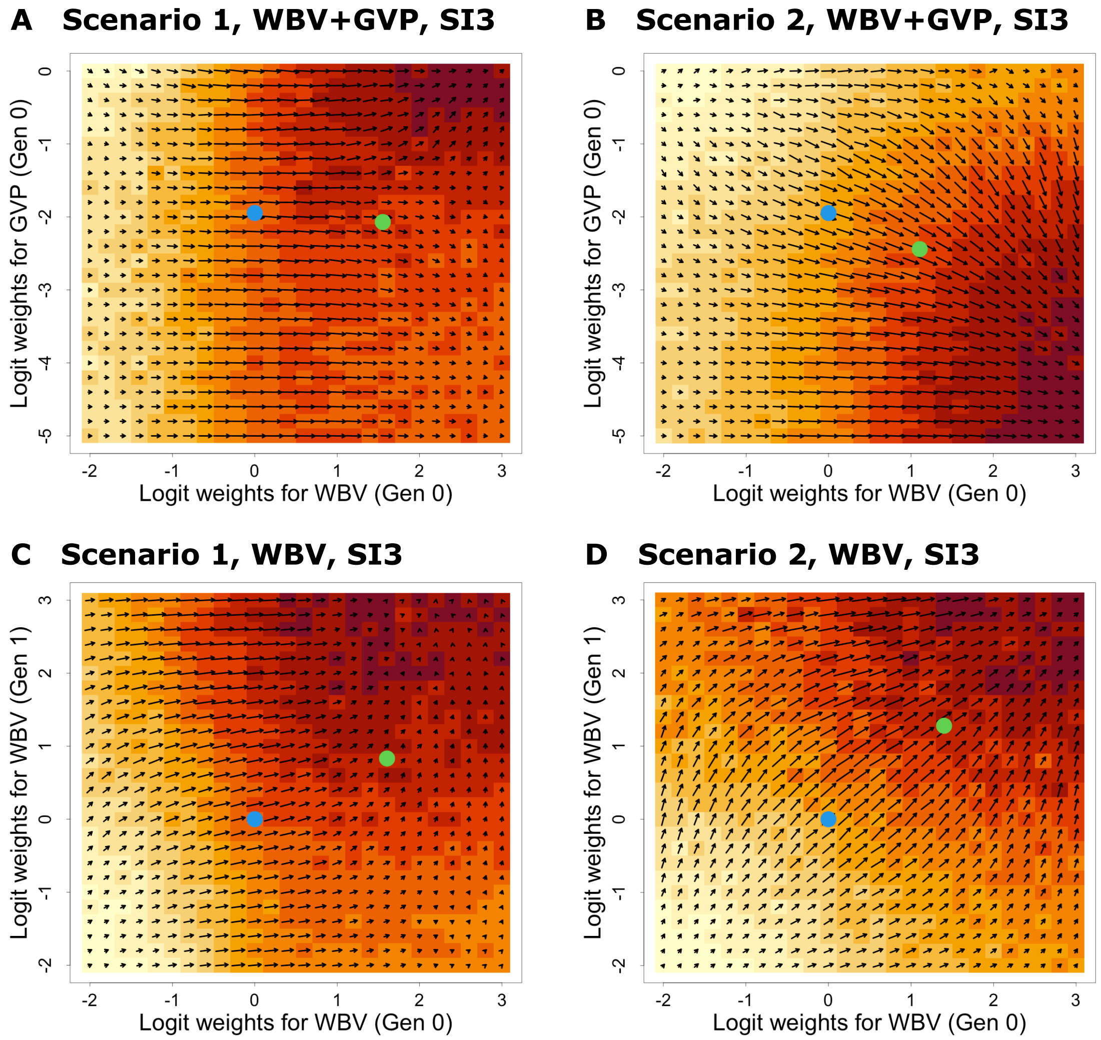


**Supplementary Figure 9. Optical flow for the gradients calculated by AD under SI3.** The function values and gradients were evaluated by AD via grid search under SI3. The horizontal and vertical axes represent the logit-transformed weighting parameters $\boldsymbol{\nu}^{(\tau)}$ in the allocation step. The background color represents the size of the function values for each set of logit-transformed weighting parameters, whereas the arrows reflect the sizes and directions of the gradients evaluated by AD. The blue and green points correspond to the initial and optimized values by the GORA2 approach. (A), (B) The gradients were evaluated for the breeding scheme with only one generation considering both WBV and GVP as candidates of $\boldsymbol{\omega}$. The horizontal and vertical axes represent the logit-transformed weighting parameters for WBV and GVP in generation 0, respectively. (C), (D) The gradients were evaluated for the breeding scheme with two generations considering only WBV as candidates of $\boldsymbol{\omega}$. The horizontal and vertical axes represent the logit-transformed weighting parameters for WBV in generations 0 and 1, respectively. (A), (C) Scenario 1. (B), (D) Scenario 2.

### Supplementary Tables

**Table S1. Optimized weighting parameters $\theta^{(\tau)}$ across different generations $\tau = 0, \ldots, T-1$ for Scenario 1 under SI2.**

| Scenario 1, SI2 | GORA1 | | GORA2 | | ORA | |
|---|---|---|---|---|---|---|
| Generation ($\tau$) | WBV | GVP | WBV | GVP | WBV | GVP |
| **0** | 3.25 | 2.71 | 3.21 | 2.40 | 1.50 | 0.17 |
| **1** | 2.44 | 2.20 | 2.50 | 0.08 | 2.83 | 0.17 |
| **2** | 2.51 | 1.80 | 2.59 | −0.02 | 2.83 | 1.5 |
| **3** | 2.26 | 1.95 | 2.47 | −0.07 | 2.83 | 0.17 |

**Table S2. Optimized weighting parameters $\theta^{(\tau)}$ across different generations $\tau = 0, \ldots, T-1$ for Scenario 1 under SI3.**

| Scenario 1, SI3 | GORA1 | | GORA2 | | ORA | |
|---|---|---|---|---|---|---|
| Generation ($\tau$) | WBV | GVP | WBV | GVP | WBV | GVP |
| **0** | 3.29 | 2.07 | 3.26 | 2.06 | 2.83 | 0.17 |
| **1** | 2.58 | 1.87 | 2.69 | 0.01 | 2.83 | 0.17 |
| **2** | 2.74 | 1.61 | 2.85 | −0.04 | 1.50 | 0.17 |
| **3** | 2.59 | 1.81 | 2.69 | −0.06 | 2.83 | 0.17 |

**Table S3. Optimized weighting parameters $\theta^{(\tau)}$ across different generations $\tau = 0, \ldots, T-1$ for Scenario 2 under SI1.**

| Scenario 2, SI1 | GORA1 | | GORA2 | | ORA | |
|---|---|---|---|---|---|---|
| Generation ($\tau$) | WBV | GVP | WBV | GVP | WBV | GVP |
| **0** | 2.36 | 0.02 | 2.44 | −0.17 | 1.06 | 0.17 |
| **1** | 2.55 | 2.20 | 2.64 | 0.56 | 0.17 | 0.17 |
| **2** | 2.32 | 2.36 | 2.63 | 0.05 | 0.17 | 0.17 |

| | | | | | | |
|---|---|---|---|---|---|---|
| **3** | 2.17 | 2.19 | 2.57 | −0.17 | 2.83 | 0.17 |

**Table S4. Optimized weighting parameters $\boldsymbol{\theta}^{(\tau)}$ across different generations $\boldsymbol{\tau} = \mathbf{0}, \dots, \boldsymbol{T} - \mathbf{1}$ for Scenario 2 under SI2.**

| **Scenario 2, SI2** | **GORA1** | | **GORA2** | | **ORA** | |
|---|---|---|---|---|---|---|
| **Generation ($\boldsymbol{\tau}$)** | **WBV** | **GVP** | **WBV** | **GVP** | **WBV** | **GVP** |
| **0** | 2.52 | 0.30 | 2.23 | −0.19 | 1.06 | 1.06 |
| **1** | 2.68 | 1.67 | 2.68 | 0.22 | 1.94 | 0.61 |
| **2** | 2.51 | 2.04 | 2.80 | −0.05 | 1.06 | 0.61 |
| **3** | 2.35 | 2.08 | 2.68 | −0.16 | 3.28 | 0.17 |

**Table S5. Optimized weighting parameters $\boldsymbol{\theta}^{(\tau)}$ across different generations $\boldsymbol{\tau} = \mathbf{0}, \dots, \boldsymbol{T} - \mathbf{1}$ for Scenario 2 under SI3.**

| **Scenario 2, SI3** | **GORA1** | | **GORA2** | | **ORA** | |
|---|---|---|---|---|---|---|
| **Generation ($\boldsymbol{\tau}$)** | **WBV** | **GVP** | **WBV** | **GVP** | **WBV** | **GVP** |
| **0** | 2.78 | 0.02 | 2.50 | −0.17 | 1.06 | 0.17 |
| **1** | 2.81 | 1.62 | 2.81 | 0.10 | 1.50 | 0.61 |
| **2** | 2.73 | 1.69 | 2.88 | −0.06 | 1.06 | 0.17 |
| **3** | 2.57 | 1.82 | 2.79 | −0.14 | 3.28 | 0.17 |

### References


Goddard,M. (2009) Genomic selection: prediction of accuracy and maximisation of long term response. *Genetica*, **136**, 245–257.

Hamazaki,K. and Iwata,H. (2024) AI-assisted selection of mating pairs through simulation-based optimized progeny allocation strategies in plant breeding. *Front. Plant Sci.*, **15**.

Jang,E. *et al.* (2017) Categorical Reparameterization with Gumbel-Softmax. In, *International Conference on Learning Representations*.

Jannink,J.-L. (2010) Dynamics of long-term genomic selection. *Genet. Sel. Evol.*, **42**, 35.

Kosambi,D.D. (1943) The estimation of map distances from recombination values. *Ann. Eugen.*, **12**, 172–175.

Liang,L. *et al.* (2007) GENOME: a rapid coalescent-based whole genome simulator. *Bioinformatics*, **23**, 1565–1567.

Maddison,C.J. *et al.* (2017) The Concrete Distribution: A Continuous Relaxation of Discrete Random Variables. In, *International Conference on Learning Representations*.

Maechler,M. *et al.* (2021) cluster: Cluster Analysis Basics and Extensions.

Müller,D. *et al.* (2017) Persistency of Prediction Accuracy and Genetic Gain in Synthetic Populations Under Recurrent Genomic Selection. *G3* , **7**, 801–811.

Müller,D. *et al.* (2018) Selection on Expected Maximum Haploid Breeding Values Can Increase Genetic Gain in Recurrent Genomic Selection. *G3* , **8**, 1173–1181.

Paszke,A. *et al.* (2019) PyTorch: An imperative style, high-performance deep learning library. *Adv. Neural Inf. Process. Syst.*, **abs/1912.01703**.

Robbins,H. and Monro,S. (1951) A Stochastic Approximation Method. *aoms*, **22**, 400–

407.

Van Inghelandt,D. *et al.* (2011) Extent and genome-wide distribution of linkage disequilibrium in commercial maize germplasm. *Theor. Appl. Genet.*, **123**, 11–20.

Yabe,S. *et al.* (2017) A simple package to script and simulate breeding schemes: The breeding scheme language. *Crop Sci.*, **57**, 1347–1354.

Zhao,H. and Speed,T.P. (1996) On genetic map functions. *Genetics*, **142**, 1369–1377.
